# Cell-type specific autophagy in root hair forming cells is essential for salt stress tolerance in *Arabidopsis thaliana*

**DOI:** 10.1101/2025.03.18.643786

**Authors:** Jierui Zhao, Christian Löfke, Kai Ching Yeung, Yixuan Chen, Yasin Dagdas

## Abstract

Autophagy is a vital cellular quality control pathway that enables plants to adapt to changing environments. By degrading damaged or unwanted components, autophagy maintains cellular homeostasis. While the organismal phenotypes of autophagy-deficient plants under stress are well-characterized, the contribution of cell-type-specific autophagy responses to whole-plant homeostasis remains poorly understood. Here, we show that root hair-forming cells (trichoblasts) of *Arabidopsis thaliana* exhibit higher autophagic flux than adjacent non-hair cells (atrichoblasts). This differential autophagy is genetically linked to cell fate determination during early development. Mutants disrupting trichoblast or atrichoblast identity lose the autophagy distinction between these cell types. Functional analyses reveal that elevated autophagy in trichoblasts is essential for sodium ion sequestration in vacuoles—a key mechanism for salt stress tolerance. Disrupting autophagy specifically in trichoblasts impairs sodium accumulation and reduces plant survival under salt stress. Conversely, cell-type-specific complementation restores both sodium sequestration and stress tolerance. Our findings uncover a cell-type-specific autophagy program in root hairs and demonstrate how developmental cues shape autophagy to enhance stress resilience. This work establishes a direct link between cell identity, autophagy, and environmental adaptation in plants.

## Introduction

Autophagy, a conserved eukaryotic degradation pathway, plays a pivotal role in maintaining cellular homeostasis by recycling damaged organelles, protein aggregates, and microbes (Gross et al., 2025). It is orchestrated by a suite of autophagy-related (ATG) proteins that mediate the *de novo* formation of a double-membraned vesicle, termed the autophagosome, which captures and delivers the cargo to the vacuole for breakdown (Avin-Wittenberg et al., 2015; Marshall & Vierstra, 2018). Autophagosome biogenesis is initiated by the ATG1 kinase complex (comprising ATG1, ATG13, ATG11, and ATG101), which integrates stress signals from upstream regulators such as TOR and SnRK1 kinase complexes to initiate autophagosome biogenesis (Mizushima, 2018; Marshall & Vierstra, 2018). Phagophore expansion depends on the phosphatidylinositol 3-kinase (PI3K) complex (VPS34, ATG6, VPS15), and membrane transfer is facilitated by proteins like ATG9 and ATG2 (Huang & Bassham, 2015). Autophagosome maturation requires the ubiquitin-like conjugation systems (ATG3, ATG4, ATG7, ATG12-ATG5-ATG16) responsible for ATG8 lipidation—a hallmark of autophagosome formation (Chung et al., 2010). While ATG proteins orchestrate autophagosome biogenesis, cargo selection and recruitment are mediated by selective autophagy receptors (SARs), such as NBR1 and ATI1 (Stolz et al., 2014; Svenning et al., 2011; Honig et al., 2012; Michaeli et al., 2014). SARs on the one hand interact with ATG8 via conserved ATG-interacting motifs (AIMs) and on the other hand contain cargo recognition domains for selective cargo recruitment (Kirkin & Rogov, 2019; Luong et al., 2022; Gross et al., 2025).

Plant autophagy is a critical adaptive mechanism enabling survival during environmental stress (Bassham et al., 2012; Avin-Wittenberg, 2019). Under abiotic stress, such as nutrient deprivation, drought, and salinity, autophagy reallocates cellular resources to sustain metabolic functions and delay senescence (Guiboileau et al., 2012; Avin-Wittenberg, 2019). The functional importance of autophagy in plants is underscored by the phenotypic consequences observed in autophagy-deficient mutants (atg mutants), which exhibit abnormal organ development, premature senescence, hypersensitivity to biotic and abiotic stresses, and metabolic imbalances (Yoshimoto et al., 2004; Minina et al., 2018; Yagyu & Yoshimoto, 2024). While organismal outcomes of autophagy defects are well-established, emerging evidence emphasizes its cell type-specific functions, allowing tailored responses across plant tissues (Feng et al., 2022). For example, in the Arabidopsis root cap, autophagy facilitates the programmed death and clearance of root cap border cells, a process critical for organized cell separation and root growth (Feng et al., 2022; Goh et al., 2022). Disruption of ATG5 in the root cap via tissue-specific CRISPR mutagenesis impairs vacuolarization and the removal of dying cells, demonstrating the developmental precision of autophagy in a spatially restricted context (Feng et al., 2022). Similarly, cell-type specific autophagy has been shown to play important roles in leaf abscission (Htwe et al., 2011; Furuta et al., 2024). However, the degree to which cell-type specific autophagy responses contribute to stress tolerance remains largely unknown.

Arabidopsis roots offer an ideal system to investigate cell type-specific autophagy as the patterning of cell types that have very different metabolic and homeostatic demands are well established. Particularly the trichoblasts and atrichoblasts, which are adjacent to each other and differentiate into root hair and non-root hair cells (Datta et al., 2011), provides an excellent system to study cell-type specific homeostatic responses. The differentiation of trichoblasts (root hair cells) and atrichoblasts (non-hair cells) in the Arabidopsis root epidermis is a complex process regulated by a combination of positional signaling, transcriptional networks, and hormonal cues (Balcerowicz et al., 2015; Salazar-Henaoet al., 2016). The fate of Arabidopsis root epidermal cells is determined by their position relative to the underlying cortical cells. Cells in contact with two cortical cells (T position) become trichoblasts, while those in contact with only one cortical cell (A position) become atrichoblasts (Balcerowicz et al., 2015; Salazar-Henaoet al., 2016). The differentiation of trichoblasts and atrichoblasts are regulated by a network of transcription factors. In A-positioned cells, the MYB transcription factor WEREWOLF (WER) forms a complex with the bHLH proteins GLABRA3 (GL3) or ENHANCER OF GLABRA3 (EGL3) and the WD40 protein TRANSPARENT TESTA GLABRA1 (TTG1) (Balcerowicz et al., 2015; Salazar-Henaoet al., 2016). This complex promotes the expression of the homeodomain protein GLABRA2 (GL2), which inhibits the expression of downstream genes related to root hair formation, leading to the atrichoblast fate (Balcerowicz et al., 2015; Salazar-Henaoet al., 2016). In T-positioned cells, the R3 MYB protein CAPRICE (CPC) (or its paralogs TRYPTYCHON [TRY] and ENHANCER OF TRY AND CPC1 [ETC1]) replaces WER in the complex, preventing GL2 expression and allowing the cells to adopt the trichoblast fate (Balcerowicz et al., 2015; Salazar-Henaoet al., 2016).

Previous studies reported autophagic flux differences between root hair cells and non-root hair cells (Guichard et al., 2024). However, the physiological and genetic basis underlying this difference remains unknown. Understanding how autophagy links with the cell type-specific transcriptional programs in Arabidopsis roots may reveal novel regulatory hubs that mediate stress adaptation and developmental plasticity.

Here, we dissected autophagic flux pattern in trichoblasts and atrichoblasts during salt stress. Using a large suite of reporters, we showed that trichoblast cells have higher autophagic flux compared the adjacent atrichoblast cells. This is encoded by the trichoblast developmental program as autophagic flux differences disappeared in genetic mutants that change cell fate. Cell-type specific CRISPR mutagenesis and complementation experiments revealed higher autophagic flux in trichoblasts is crucial for salt stress tolerance. Altogether, by mapping autophagy dynamics at cellular resolution, we uncovered a cell-type specific autophagy response that is crucial for stress resilience.

## Results

### Trichoblasts exhibit significantly higher autophagic flux compared to atrichoblasts in the root maturation zone of *Arabidopsis thaliana*

Trichoblasts and atrichoblasts are organized in a highly structured adjacent manner. To investigate whether these two cell types exhibit distinct autophagic activities, we first measured the autophagic flux in trichoblasts and atrichoblasts at the root maturation zone of Arabidopsis wildtype Col-0 expressing GFP-ATG8A, under control and two autophagy-inducing conditions (NaCl stress and nitrogen starvation), using confocal microscopy. To accurately identify trichoblasts and atrichoblasts in our live-cell imaging experiments, we adhered to the following criteria: an epidermal cell was classified as a trichoblast if it was adjacent to two cortical cells, whereas it was classified as an atrichoblast if it was adjacent to only one cortical cell (Figure S1). Under both control and autophagy-inducing conditions, confocal microscopy revealed that trichoblasts exhibit significantly higher autophagic flux compared to the adjacent atrichoblast cells (Figure 1 and S2). Similar to ATG8A, the other eight ATG8 isoforms (ATG8B to I) also exhibited higher autophagic flux in trichoblast cells compared to atrichoblasts at the root maturation zone under control, NaCl stress, and nitrogen-starvation conditions (Figure 1 and S2). Collectively, these results demonstrate that trichoblasts in the root maturation zone exhibit significantly higher autophagic activity compared to the adjacent atrichoblast cells. This observation prompted us to further explore this cell type specific autophagic response in more detail.

**Figure 1.**
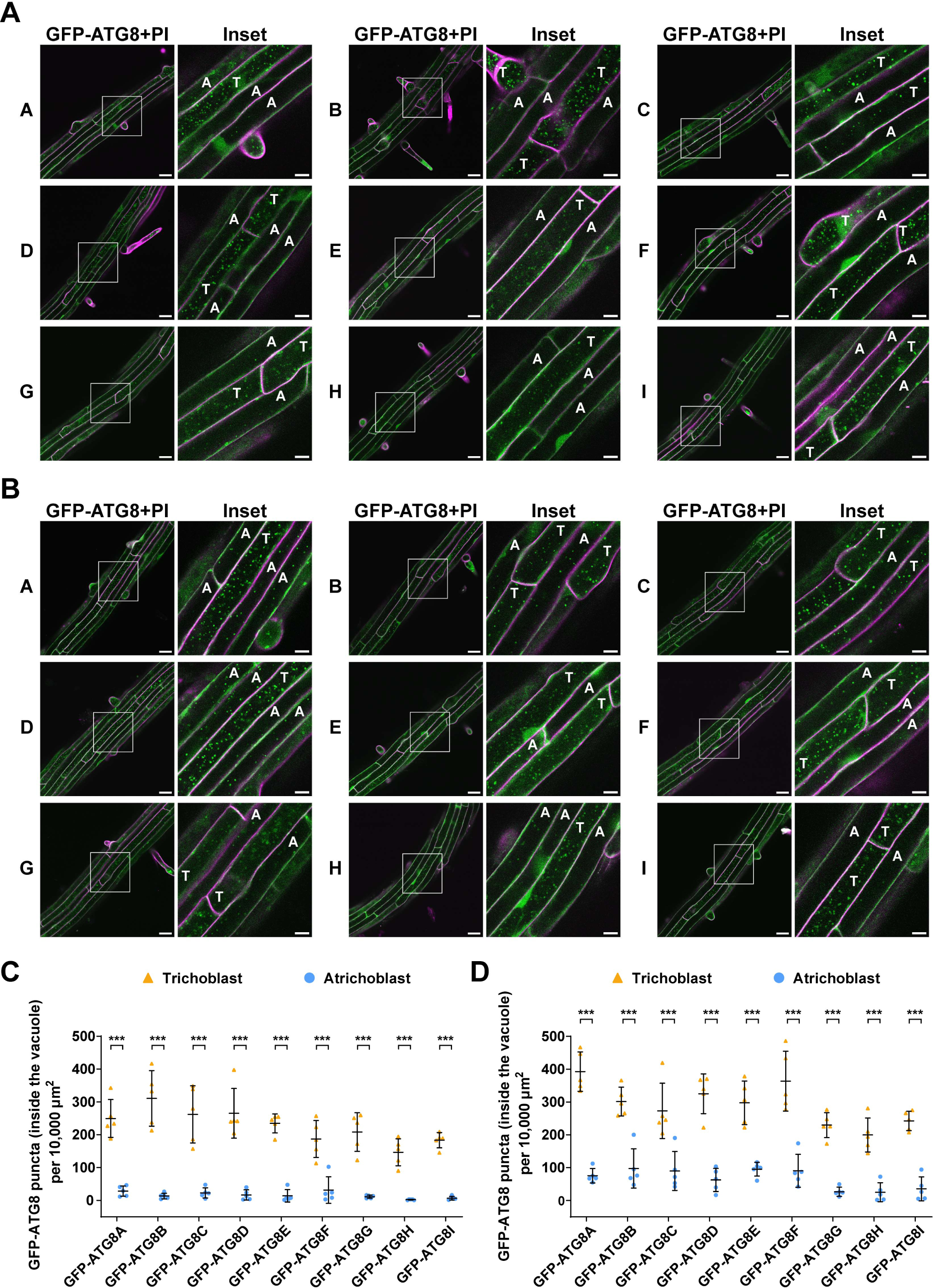
Trichoblasts exhibit significantly higher autophagic flux than neighboring atrichoblasts under both control and NaCl treatments in *Arabidopsis thaliana* root maturation zone. **(A)** Confocal microscopy images of the trichoblasts and atrichoblasts at root maturation zone of Arabidopsis wildtype Col-0 expressing ProUBQ10:GFP-ATG8A–I isoforms under control treatment. 5-d-old Arabidopsis seedlings were incubated in control 1/2 MS media containing 2 μm concanamycin A for 2 h before imaging. Representative images of 5 replicates are shown. Area highlighted in the white-boxed region in the GFP-ATG8+PI panel was further enlarged and presented in the inset panel. Scale bars, 30 μm. Inset scale bars, 10 μm. Green color, GFP-ATG8A–I isoforms. Magenta color, propidium iodide dye. T, trichoblasts. A, atrichoblasts. **(B)** Confocal microscopy images of the trichoblasts and atrichoblasts at root maturation zone of Arabidopsis wildtype Col-0 expressing ProUBQ10:GFP-ATG8A–I isoforms under NaCl treatment. 5-d-old Arabidopsis seedlings were incubated in 1/2 MS media containing 50 mM NaCl + 1 μm concanamycin A for 40-60 min before imaging. Representative images of 5 replicates are shown. Area highlighted in the white-boxed region in the GFP-ATG8+PI panel was further enlarged and presented in the inset panel. Scale bars, 30 μm. Inset scale bars, 10 μm. Green color, GFP-ATG8A–I isoforms. Magenta color, propidium iodide dye. T, trichoblasts. A, atrichoblasts. **(C)** Quantification of the GFP-ATG8 puncta inside the vacuole per normalized area (10,000 μm^2^) of the trichoblasts and atrichoblasts imaged in A. Bars indicate the mean ± SD of 5 replicates. Two-tailed and paired student t-tests were performed to analyze the significance of GFP-ATG8 puncta density differences between the trichoblasts and the atrichoblasts. ***, P < 0.001. **(D)** Quantification of the GFP-ATG8 puncta inside the vacuole per normalized area (10,000 μm^2^) of the trichoblasts and atrichoblasts imaged in B. Bars indicate the mean ± SD of 5 replicates. Two-tailed and paired student t-tests were performed to analyze the significance of GFP-ATG8 puncta density differences between the trichoblasts and the atrichoblasts. ***, P < 0.001.

### Genetic basis of the higher autophagic flux in trichoblast cells

The development of trichoblasts and atrichoblasts is regulated by a well-characterized genetic program (Balcerowicz et al., 2015; Salazar-Henao et al., 2016). While the formation of root hairs from trichoblasts at the maturation zone results in obvious surface differences, the determination of cell type identity occurs at the meristematic zone (Löfke et al., 2013). Given that both root hair structure and cell type identity could contribute to the observed differences in autophagic activity, we investigated autophagic activity in a series of Arabidopsis root epidermal development mutants. We selected four representative mutants that were previously shown to affect cell fate and identity: *rhd6 rsl1*, *cpc try*, *gl2*, and *wer myb23* (Figure 2A). *rhd6 rsl1* exhibits ectopic non-hair cells at trichoblast positions, while *gl2* displays ectopic hair cells at atrichoblast positions. However, both *rhd6 rsl1* and *gl2* retain meristematic cell type determination, as evidenced by meristematic vacuolar biogenesis phenotype (Löfke et al., 2013). In contrast, *cpc try* and *wer myb23* lose cell type determination at the meristematic stage, despite *cpc try* having ectopic non-hair cells and *wer myb23* having ectopic hair cells (Figure 2A). To test the genetic basis of higher autophagic flux in trichoblast cells, we expressed GFP-ATG8A in these four mutants, as well as in *atg5* as an autophagy-defective control, and performed live-cell imaging experiments similar to those described in Figure 1. Analysis of autophagic flux in epidermal cells at trichoblast and atrichoblast positions revealed that *rhd6 rsl1* and *gl2* maintained the autophagic flux difference, whereas *cpc try* and *wer myb23* lost this difference. This suggests cell fate defined at the meristematic zone underlies the differential autophagic activity between trichoblasts and atrichoblasts (Figure 2B–2E and S3). However, the presence of root hair structures also appeared to slightly enhance autophagic flux, as evidenced by the slightly lower autophagic flux in trichoblast-positioned cells of *rhd6 rsl1* compared to Col-0 and *gl2*, and the slightly higher autophagic flux in atrichoblast-positioned cells of *gl2* compared to other lines (Figure 2B–2E and S3).

**Figure 2.**
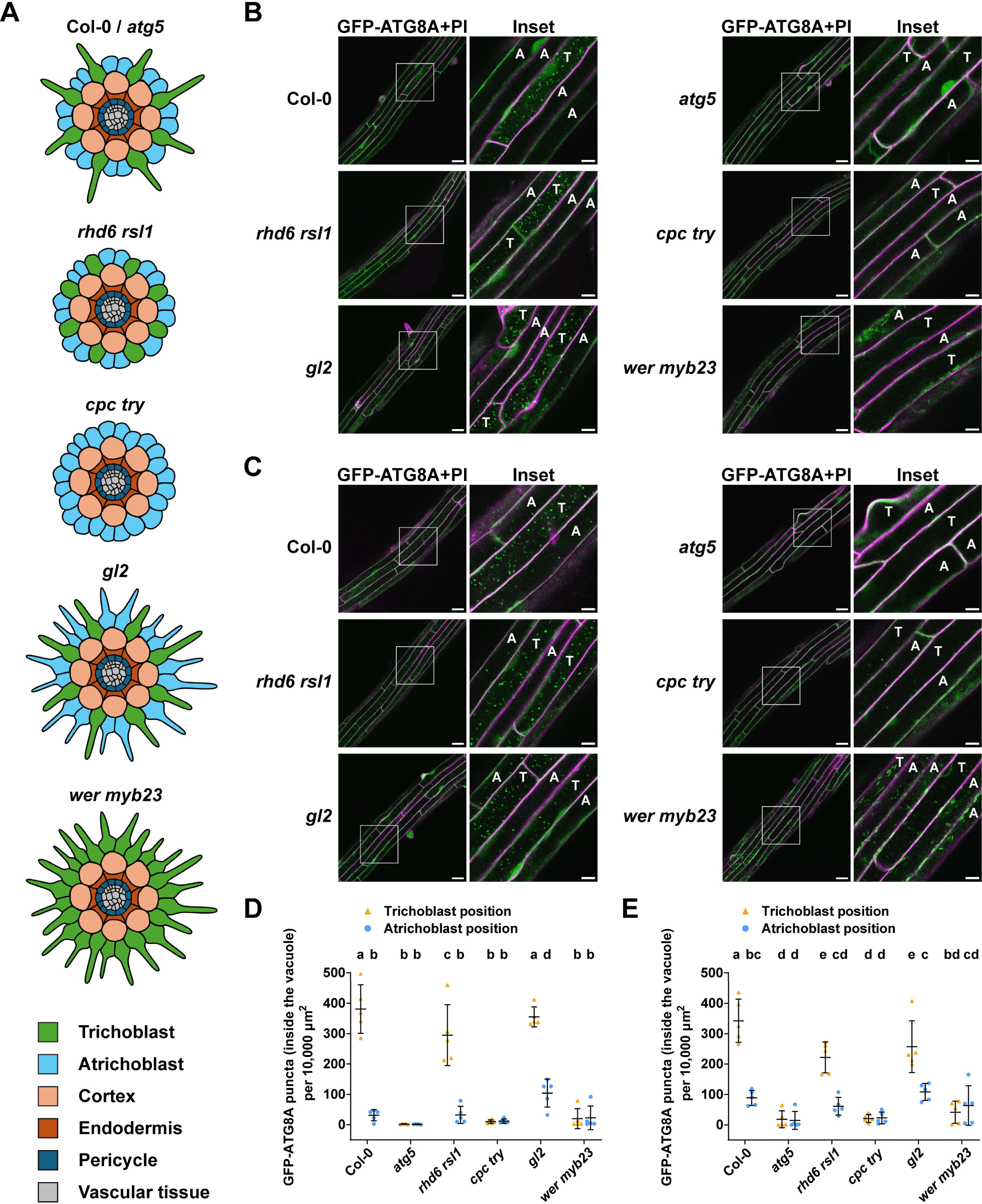
Genetic basis of the autophagic flux difference between trichoblasts and atrichoblasts. **(A)** Schematic diagram showing the trichoblast-atrichoblast distribution pattern in Arabidopsis wildtype Col-0, the autophagy-defective mutant *atg5* and different root-hair development mutant lines *rhd6 rsl1*, *cpc try*, *gl2* and *wer myb23*. **(B)** Confocal microscopy images of epidermal cells at the root maturation zone of Col-0, *atg5*, *rhd6 rsl1*, *cpc try*, *gl2* and *wer myb23* expressing ProUBQ10:GFP-ATG8A under control treatment. 5-d-old Arabidopsis seedlings were incubated in control 1/2 MS media containing 2 μm concanamycin A for 2 h before imaging. Representative images of 5 replicates are shown. Area highlighted in the white-boxed region in the GFP-ATG8A+PI panel was further enlarged and presented in the inset panel. Scale bars, 30 μm. Inset scale bars, 10 μm. Green color, GFP-ATG8A. Magenta color, propidium iodide dye. Note, T indicates the trichoblast positions (adjacent to two cortex cells) and A indicates the atrichoblast positions (adjacent to only one cortex cell), rather than the cell identities as mutants are affected in cell identity development. **(C)** Confocal microscopy images of epidermal cells at the root maturation zone of Col-0, *atg5*, *rhd6 rsl1*, *cpc try*, *gl2* and *wer myb23* expressing ProUBQ10:GFP-ATG8A under NaCl treatment. 5-d-old Arabidopsis seedlings were incubated in 1/2 MS media containing 50 mM NaCl + 1 μm concanamycin A for 40-60 min before imaging. Representative images of 5 replicates are shown. Area highlighted in the white-boxed region in the GFP-ATG8A+PI panel was further enlarged and presented in the inset panel. Scale bars, 30 μm. Inset scale bars, 10 μm. Green color, GFP-ATG8A. Magenta color, propidium iodide dye. Note, T indicates the trichoblast positions (adjacent to two cortex cells) and A indicates the atrichoblast positions (adjacent to only one cortex cell), rather than the cell identities as mutants are affected in cell identity development. **(D)** Quantification of the GFP-ATG8A puncta inside the vacuole per normalized area (10,000 μm^2^) of the cells at the trichoblast positions and the atrichoblast positions imaged in B. Bars indicate the mean ± SD of 5 replicates. Paired repeated measures one-way ANOVA and Fisher’s LSD tests were used to analyze the differences of the number of the GFP-ATG8A puncta between each group. Family-wise significance and confidence level, 0.05 (95% confidence interval). **(E)** Quantification of the GFP-ATG8A puncta inside the vacuole per normalized area (10,000 μm^2^) of the cells at the trichoblast positions and the atrichoblast positions imaged in C. Bars indicate the mean ± SD of 5 replicates. Paired repeated measures one-way ANOVA and Fisher’s LSD tests were used to analyze the differences of the number of the GFP-ATG8A puncta between each group. Family-wise significance and confidence level, 0.05 (95% confidence interval).

Notably, *wer myb23* exhibited an unexpected autophagic flux pattern, with all root epidermal cells displaying very low flux (Figure 2B–2E and S3). Strikingly, GFP-ATG8A signals in *wer myb23* were predominantly localized to the endoplasmic reticulum (ER) and ER bodies under both control and autophagy-inducing conditions (Figure S4A). Colocalization experiments in Col-0 and *wer myb23* that co-express GFP-ATG8A with the ER marker DDRGK1-mCherry (Gerakis et al., 2019) confirmed the entrapment of GFP-ATG8A signal at the ER (Figure S4B). These findings suggest the transcriptional network that underlies root epidermal cell development impinges on autophagosome biogenesis and cell-type identity dictates the higher autophagic flux in trichoblast cells.

### Establishment of cell-type specific CRISPR mutagenesis and complementation lines to investigate the physiological relevance of higher autophagic flux in trichoblast cells

To further investigate the physiological significance of the differential autophagic activity between trichoblasts and atrichoblasts, we established two genetic approaches to modulate autophagic flux in trichoblasts. First, using trichoblast-specific promoters *ProEXP7* and *ProRHD6* (Figure S5) and tissue-specific CRISPR knockout (CRISPR-TSKO) technology, we aimed to disrupt autophagic flux specifically in trichoblasts. We expressed *ProEXP7*-driven and *ProRHD6*-driven Cas9 to knockout *ATG5* in Col-0 expressing ProUBQ10:mCherry-ATG8E (mCherry-ATG8E in Col-0), generating the lines TSKO-E (*ProEXP7*-driven) and TSKO-R (*ProRHD6-*driven) (Figure 3A). Interestingly, under both control and NaCl stress conditions, only TSKO-R exhibited a significant reduction in mCherry-ATG8E-marked autophagic flux in trichoblasts at the maturation zone, while TSKO-E maintained autophagic flux levels similar to Col-0 (Figure S6). Given that *ProEXP7* is expressed exclusively in trichoblasts at the maturation zone, whereas *ProRHD6* is also expressed in trichoblasts at the meristematic and elongation zones (Figure S5), we hypothesized that pre-existing ATG5 is stable enough to reach maturation zone cells. Additionally, we confirmed that TSKO-R did not exhibit reduced autophagic flux in cells that don’t express *RHD6*, such as those in the stele and cotyledon epidermis (Figure S7), indicating that autophagy remained intact in non-*ProRHD6* expressing regions. Based on these results, we selected TSKO-R for further studies.

**Figure 3.**
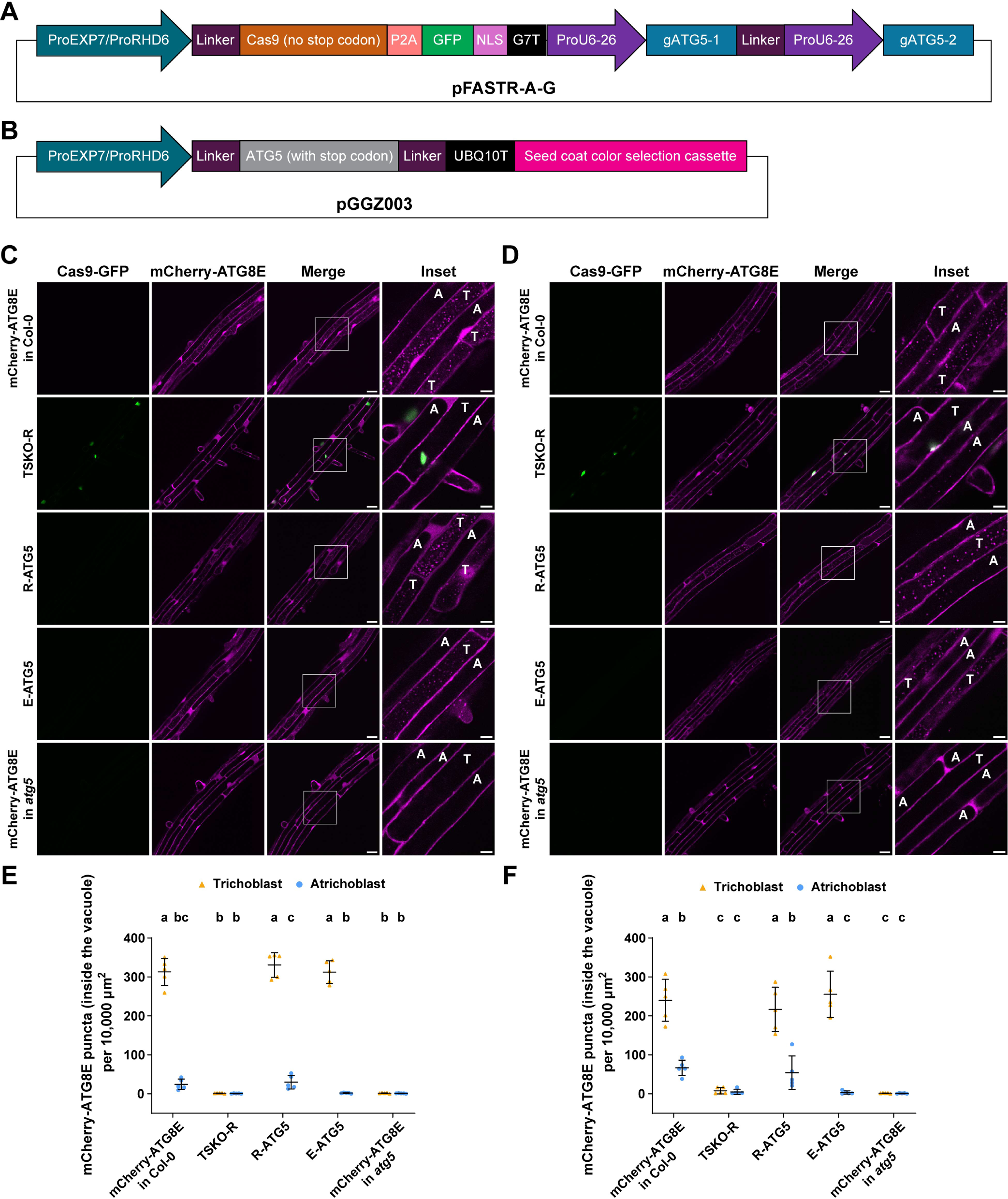
Tissue specific CRISPR mutagenesis and complementation of *ATG5* confirm higher autophagic flux in trichoblast cells. **(A)** Schematic diagram showing the design of the trichoblast-specific *ATG5* CRISPR mutagenesis plasmid expressed in wild type Col-0 expressing ProUBQ10:mCherry-ATG8E(mCherry-ATG8E in Col-0). **(B)** Schematic diagram showing the design of the trichoblast-specific *ATG5* complementation plasmid transformed to complement the autophagy-defective mutant *atg5* rexpressing ProUBQ10:mCherry-ATG8E (mCherry-ATG8E in *atg5*). **(C)** Confocal microscopy images of trichoblasts and atrichoblasts of Arabidopsis lines mCherry-ATG8E in Col-0, TSKO-R (*ATG5* mutagenized using *ProRHD6*-driven Cas9), R-ATG5 (*atg5* mutant complemented with *ProRHD6*-driven *ATG5*), E-ATG5 (*atg5* mutant complemented with *ProEXP7*-driven *ATG5*) and mCherry-ATG8E in *atg5* under control treatment. 5-d-old Arabidopsis seedlings were incubated in control 1/2 MS media containing 2 μm concanamycin A for 2 h before imaging. Representative images of 5 replicates are shown. Area highlighted in the white-boxed region in the merge panel was further enlarged and presented in the inset panel. Scale bars, 30 μm. Inset scale bars, 10 μm. T, trichoblasts. A, atrichoblasts. **(D)** Confocal microscopy images of trichoblasts and atrichoblasts of Arabidopsis lines mCherry-ATG8E in Col-0, TSKO-R (*ATG5* mutagenized using *ProRHD6*-driven Cas9), R-ATG5 (*atg5* mutant complemented with *ProRHD6*-driven *ATG5*), E-ATG5 (*atg5* mutant complemented with *ProEXP7*-driven *ATG5*) and mCherry-ATG8E in *atg5* under NaCl stress treatment. 5-d-old Arabidopsis seedlings were incubated in 1/2 MS media containing 50 mM NaCl + 1 μm concanamycin A for 1 h before imaging. Representative images of 5 replicates are shown. Area highlighted in the white-boxed region in the merge panel was further enlarged and presented in the inset panel. Scale bars, 30 μm. Inset scale bars, 10 μm. T, trichoblasts. A, atrichoblasts. **(E)** Quantification of the mCherry-ATG8E puncta inside the vacuole per normalized area (10,000 μm^2^) of the trichoblasts and atrichoblasts imaged in C. Bars indicate the mean ± SD of 5 replicates. Paired repeated measures one-way ANOVA and Fisher’s LSD tests were used to analyze the differences of the number of the mCherry-ATG8E puncta between each group. Family-wise significance and confidence level, 0.05 (95% confidence interval). **(F)** Quantification of the mCherry-ATG8E puncta inside the vacuole per normalized area (10,000 μm^2^) of the trichoblasts and atrichoblasts imaged in D. Bars indicate the mean ± SD of 5 replicates. Paired repeated measures one-way ANOVA and Fisher’s LSD tests were used to analyze the differences of the number of the mCherry-ATG8E puncta between each group. Family-wise significance and confidence level, 0.05 (95% confidence interval).

In parallel, we generated Arabidopsis lines in which autophagy was specifically rescued in trichoblasts. We expressed *ProEXP7*-driven and *ProRHD6*-driven *ATG5* in Arabidopsis *atg5* mutant expressing ProUBQ10:mCherry-ATG8E (mCherry-ATG8E in *atg5*), creating the complementation lines R-ATG5 (*ProRHD6*-driven) and E-ATG5 (*ProEXP7*-driven) (Figure 3B). Subsequently, we compared mCherry-ATG8E-marked autophagic flux in Col-0, TSKO-R, R-ATG5, E-ATG5, and *atg5* under control and NaCl stress conditions. TSKO-R lost the autophagic flux difference between trichoblasts and atrichoblasts, resembling *atg5* mutant. In contrast, both R-ATG5 and E-ATG5 restored the autophagic flux difference to levels comparable to mCherry-ATG8E in Col-0 (Figure 3C–3F). These results demonstrate that TSKO-R, R-ATG5, and E-ATG5 allow us to study the physiological relevance of higher autophagic flux in trichoblast cells.

### Higher autophagic flux in trichoblasts is necessary and sufficient to compartmentalize sodium ions in trichoblast vacuoles, thereby contributing to salt stress tolerance

Next, we set out to explore the physiological relevance of higher flux in trichoblast cells. Previous studies have shown that autophagy was required for sodium ion accumulation in the vacuole of root meristem cells (Luo et al., 2017). This finding inspired us to check if higher flux in trichoblast cells is correlated with the accumulation of sodium in trichoblast vacuoles. First, we measured sodium ion concentrations in the vacuoles of epidermal cells at the root maturation zone in Col-0 and three autophagy-deficient mutants (*atg5*, *atg8*, and *atg16*), using the CoroNa Green AM staining method (Park et al., 2009; Luo et al, 2017). Notably, only trichoblasts in Col-0 exhibited significantly higher sodium ion concentrations compared to atrichoblasts (CoroNa Green AM fluorescence ratios > 1.5), whereas all autophagy-defective mutants lost this difference (CoroNa Green AM fluorescence ratios around 1) (Figure 4A and 4B). This prompted us to hypothesize that sodium accumulation in trichoblast vacuoles could help plants tolerate NaCl stress. To test our hypothesis, we grew seeds of Col-0 and autophagy-deficient mutant lines on normal 1/2 MS media for 6 days, then transferred them to 1/2 MS media supplemented with 150 mM NaCl for 4 days. Col-0 seedlings displayed significantly higher survival rates (measured as the percentage of non-etiolated leaves) compared to the autophagy-deficient lines (Figure 4C and 4D), supporting our hypothesis.

**Figure 4.**
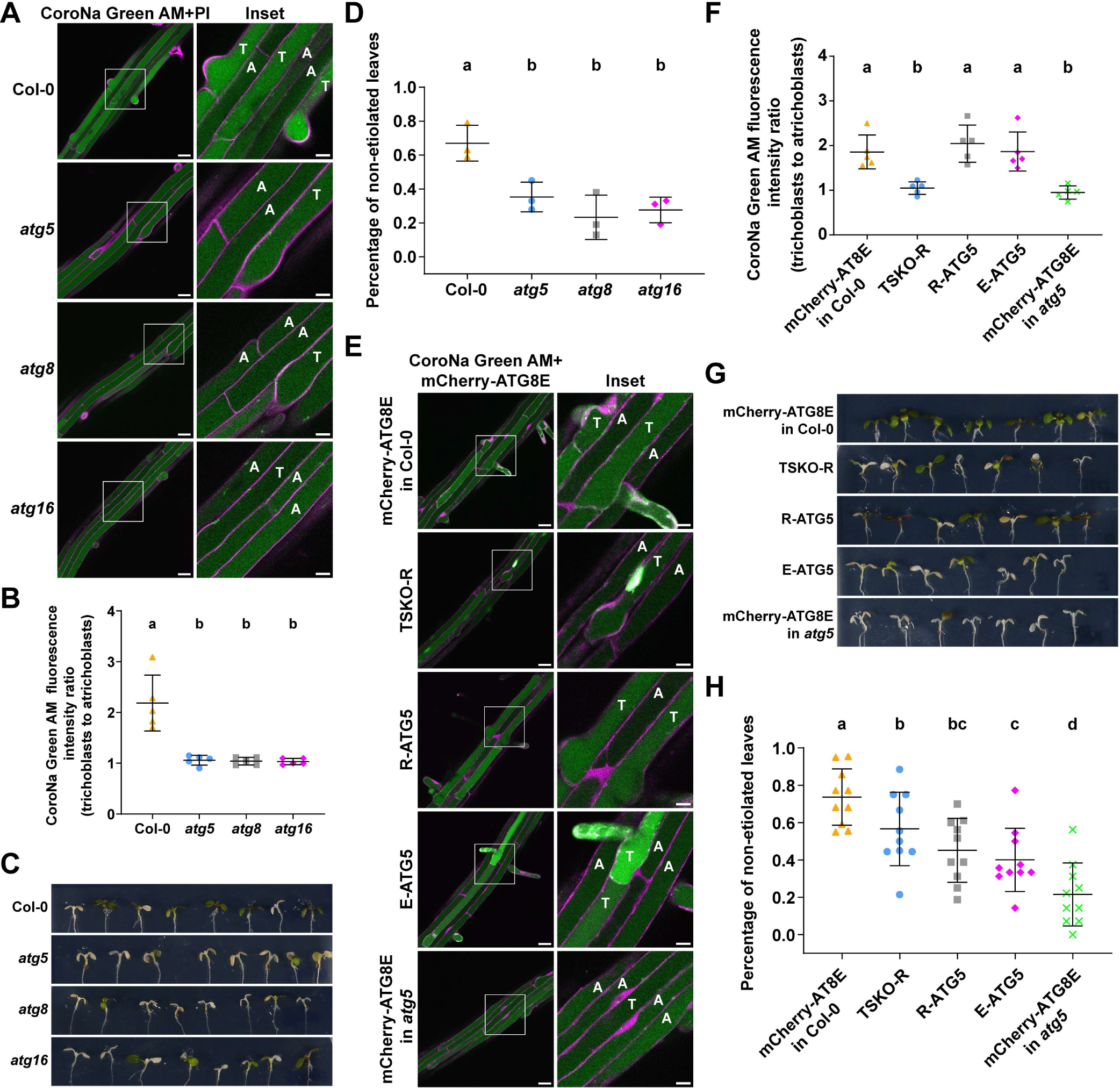
High autophagic flux in trichoblasts regulates sodium accumulation in the vacuole and is essential for coping with NaCl stress. **(A)** Confocal microscopy images showing the sodium ion accumulation in the vacuoles of trichoblasts and atrichoblasts at the root maturation zone of Arabidopsis wildtype Col-0 and the autophagy-defective mutants *atg5*, *atg8* and *atg16* indicated by CoroNa Green AM staining. 5-d-old Arabidopsis seedlings were incubated in control 1/2 MS media for 30 min and were subsequently incubated in control 1/2 MS media containing 2 μm CoroNa Green AM for 30 min before imaging. Representative images of 5 replicates are shown. Area highlighted in the white-boxed region in the CoroNa Green+PI panel was further enlarged and presented in the inset panel. Scale bars, 30 μm. Inset scale bars, 10 μm. Green color, CoroNa Green AM sodium indicator. Magenta color, propidium iodide dye. T, trichoblasts. A, atrichoblasts. **(B)** Quantitative analysis of the vacuolar CoroNa Green AM fluorescence intensity ratio between trichoblasts and atrichoblasts of the Arabidopsis seedlings imaged in A. Bars indicate the mean ± SD of 5 replicates. Paired repeated measures one-way ANOVA and Fisher’s LSD tests were performed to analyze the significance of the CoroNa Green AM fluorescence intensity ratio differences between each group. Family-wise significance and confidence level, 0.05 (95% confidence interval). **(C)** Phenotypic characterization of the seedlings of Arabidopsis wildtype Col-0 and the autophagy-defective mutants *atg5*, *atg8* and *atg16* under NaCl stress treatment. Arabidopsis seeds were vertically grown on 1/2 MS media plates (+1% plant agar) for 6 d and the 6-d-old seedlings were subsequently transferred to 1/2 MS media plates (+1% plant agar) containing 150 mM NaCl and vertically grown for 4 d. Plants were grown at 21°C under LEDs with 70 μM/m^2^/s and a 14 h light/10 h dark photoperiod. Representative images of 3 replicates are shown. **(D)** Quantitative analysis of the percentage of non-etiolated leaves of Arabidopsis seedlings imaged in C. Bars indicate the mean ± SD of 3 replicates. Paired repeated measures one-way ANOVA and Fisher’s LSD tests were performed to analyze the significance of the percentage differences between each group. Family-wise significance and confidence level, 0.05 (95% confidence interval). **(E)** Confocal microscopy images showing the sodium ion concentrations in the vacuoles of trichoblasts and atrichoblasts at the root maturation zone of Arabidopsis lines mCherry-ATG8E in Col-0, TSKO-R (*ATG5* mutagenized using *ProRHD6*-driven Cas9), R-ATG5 (*atg5* mutant complemented with *ProRHD6*-driven *ATG5*), E-ATG5 (*atg5* mutant complemented with *ProEXP7*-driven *ATG5*) and mCherry-ATG8E in *atg5* indicated by CoroNa Green AM staining. 5-d-old Arabidopsis seedlings were incubated in control 1/2 MS media for 30 min and were subsequently incubated in control 1/2 MS media containing 2 μm CoroNa Green AM for 30 min before imaging. Representative images of 5 replicates are shown. Area highlighted in the white-boxed region in the CoroNa Green AM+mCherry-ATG8E panel was further enlarged and presented in the inset panel. Scale bars, 30 μm. Inset scale bars, 10 μm. Green color, CoroNa Green AM sodium indicator (and the nuclear signals of Cas9-GFP in TSKO-R). Magenta color, mCherry-ATG8E. T, trichoblasts. A, atrichoblasts. **(F)** Quantitative analysis of the vacuolar CoroNa Green AM fluorescence intensity ratio between trichoblasts and atrichoblasts of the Arabidopsis seedlings imaged in E. Bars indicate the mean ± SD of 5 replicates. Paired repeated measures one-way ANOVA and Fisher’s LSD tests were performed to analyze the significance of the CoroNa Green AM fluorescence intensity ratio differences between each group. Family-wise significance and confidence level, 0.05 (95% confidence interval). **(G)** Phenotypic characterization of the seedlings of Arabidopsis lines mCherry-ATG8E in Col-0, TSKO-R (*ATG5* mutagenized using *ProRHD6*-driven Cas9), R-ATG5 (*atg5* mutant complemented with *ProRHD6*-driven *ATG5*), E-ATG5 (*atg5* mutant complemented with *ProEXP7*-driven *ATG5*) and mCherry-ATG8E in *atg5* under NaCl stress treatment. Arabidopsis seeds were vertically grown on 1/2 MS media plates (+1% plant agar) for 6 d and the 6-d-old seedlings were subsequently transferred to 1/2 MS media plates (+1% plant agar) containing 150 mM NaCl and vertically grown for 4 d. Plants were grown at 21°C under LEDs with 70 μM/m^2^/s and a 14 h light/10 h dark photoperiod. Representative images of 10 replicates are shown. **(H)** Quantitative analysis of the percentage of non-etiolated leaves of Arabidopsis seedlings imaged in G. Bars indicate the mean ± SD of 10 replicates. Paired repeated measures one-way ANOVA and Fisher’s LSD tests were performed to analyze the significance of the percentage differences between each group. Family-wise significance and confidence level, 0.05 (95% confidence interval).

Next, we tested if higher flux in trichoblast cells contribute to NaCl stress tolerance, using the tools we established. CoroNa Green AM staining results revealed that TSKO-R, which lacks autophagic flux differences between trichoblasts and atrichoblasts, also lost the sodium ion concentration difference (Figure 4E and 4F). Conversely, the complementation lines R-ATG5 and E-ATG5, which restored autophagic flux differences, also re-established the sodium ion concentration difference between trichoblasts and atrichoblasts (Figure 4E and 4F). Given that only a subset of cells in TSKO-R lost autophagy and only a subset in R-ATG5 and E-ATG5 regained autophagy, we predicted that these lines would exhibit intermediate survival rates between Col-0 and *atg5*. Indeed, phenotyping assays using the same approach as described above demonstrated significantly lower survival rate for TSKO-R line and significantly higher survival rates for R-ATG5 and E-ATG5 lines (Figure 4G and 4H).

Collectively, these results demonstrate that autophagy mediates sodium ion sequestration in trichoblast vacuoles, thereby contributing to NaCl stress tolerance in Arabidopsis seedlings. The precise mechanism underlying this process remains to be elucidated, but our findings highlight the importance of trichoblast-specific autophagy in salt stress responses.

## Discussion

Our study uncovers a critical role for cell-type-specific autophagy in trichoblasts, linking higher autophagic flux to sodium ion sequestration and salt stress tolerance in *Arabidopsis thaliana*. Under both control and stress conditions, trichoblasts exhibit significantly higher autophagic activity than atrichoblasts and this difference is essential for vacuolar sodium accumulation and plant survival under salinity.

A recent study has demonstrated that under salt stress conditions, Arabidopsis transports sodium ions to the vacuole to prevent toxicity. This compartmentalization is mediated by SOS1-transporter that is localized at the tonoplast (Ramakrishna et al., 2025). Our findings suggest elevated autophagic flux in trichoblasts correlates with enhanced capacity to sequester sodium ions in vacuoles. We envision two scenarios that may explain autophagy-mediated sodium sequestration in the vacuole: (1) Autophagy could support sodium compartmentalization via modulating the turnover or activity of sodium transporters. Previous studies in mammalian cells have shown that ion channels could modulate autophagy by ion fluxes in and out of lysosomes and they themselves could be modulated by autophagic recycling (Kondratskyi et al., 2018). It will be interesting to test if NHX family transporters, including SOS1 are regulated by autophagic turnover. (2) Alternatively, metalloproteins that can carry sodium ions in bulk could be delivered to the vacuole via autophagy. This is akin to ferritinophagy that is mediated by the NCOA4 selective autophagy receptor (Mancias et al., 2014). NCOA4 selectively binds ferritin and delivers ferritin molecules to the lysosomes to release iron (Wang et al., 2023). Similar to ferritinophagy, sodium binding metalloproteins could be targeted by autophagy to rapidly deliver sodium ions to the vacuole and prevent cytotoxicity. These two scenarios focus on degradative function of autophagy. Alternatively, higher autophagic flux could improve cellular homeostasis, enabling trichoblasts to maintain ion balance more effectively under stress. Future studies that investigate autophagic cargo under salt stress conditions could reveal which of these scenarios underlie elevated autophagic flux in trichoblasts.

The differential autophagic flux between trichoblasts and atrichoblasts originates during cell fate specification at the meristematic zone, as evidenced by epidermal mutants (Figure 2 and S3). Mutants disrupting early trichoblast/atrichoblast (T/A) identity (*cpc try*, *wer myb23*) abolished autophagic differences, while those retaining cell fate (*rhd6 rsl1*, *gl2*) preserved them. This suggests that developmental programs dictating T/A identity directly regulate autophagy. Transcription factors or signaling molecules common to both pathways may mediate this crosstalk. The striking mislocalization of ATG8A to the ER in *wer myb23* mutants further highlights this interplay, implying that cell fate regulators impinge on the autophagosome biogenesis machinery. This unexpected finding opens new avenues to explore how developmental signalling could spatially regulate autophagy.

Our findings underscore the importance of studying autophagy at cellular resolution. Whole-organism approaches may overlook critical tissue- or cell-type-specific dynamics, as demonstrated by the stark contrast between trichoblasts and atrichoblasts. Readily available high-resolution tools such as CRISPR-TSKO, single-cell omics, and live cell imaging allows us to move from whole organism to cell-type specific autophagy studies, and opens the door to address a fundamental yet unanswered question in cell biology: How autophagy responses in different cell types are coordinated to establish an organismal homeostatic response?

By manipulating autophagy specifically in trichoblasts using CRISPR-TSKO and ATG5 complementation lines, we demonstrated its necessity and sufficiency for sodium accumulation and salt tolerance. Disrupting autophagy in trichoblasts (TSKO-R) abolished sodium differences and reduced survival under salt stress, while restoring it (R-ATG5, E-ATG5) reinstated both. These results suggest that enhancing autophagy in stress-responsive cell types, such as trichoblasts, could improve crop resilience without compromising growth. For agriculture, this strategy could mitigate yield losses in saline soils, particularly in staple crops where root hair function is pivotal for nutrient uptake.

## Materials and methods

### Plant material and cloning procedure

All *Arabidopsis thaliana* lines used in this study are listed in Table 1. All the plasmids established in this study are listed in Table 2. The primers used for genotyping and cloning are listed in Table 3. Synthetic sequences for plasmid construction were also listed in Table 3.

**Table 1.**
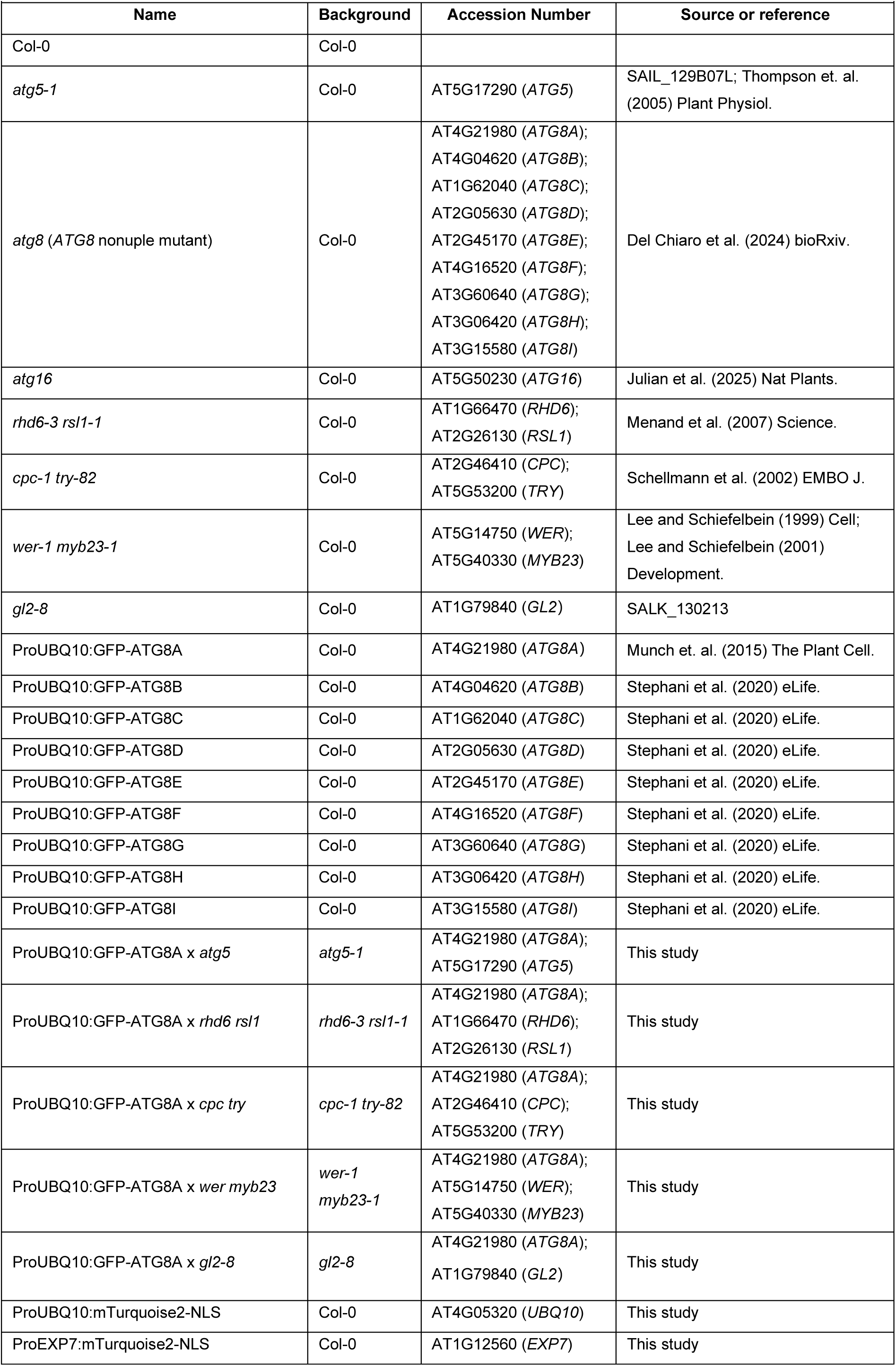
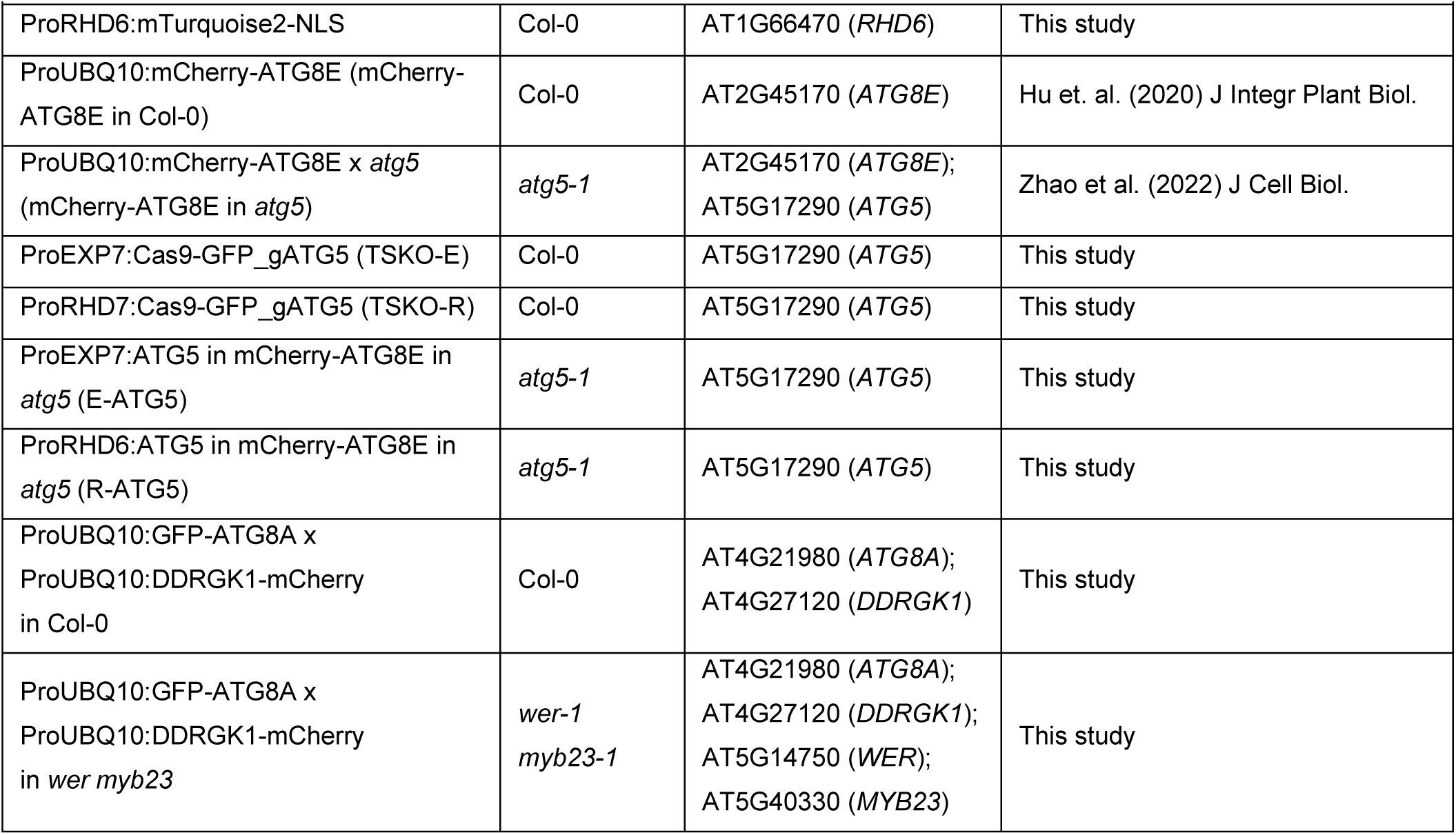
*Arabidopsis thaliana* lines used in this study.

**Table 2.** List of plasmids used in this study.

| Name | Accession number | Expression System | Backbone | Source or reference |
| --- | --- | --- | --- | --- |
| pGG-A-ProEXP7-B | AT1G12560 (*EXP7*) | *Escherichia coli* | pGGA000 | This study |
| pGG-A-ProRHD6-B | AT1G66470 (*RHD6*) | *Escherichia coli* | pGGA000 | This study |
| ProUBQ10:mTurquoise2-NLS | AT4G05320 (*UBQ10*) | *Arabidopsis thaliana* | pGGZ003 | This study |
| ProEXP7:mTurquoise2-NLS | AT1G12560 (*EXP7*) | *Arabidopsis thaliana* | pGGZ003 | This study |
| ProRHD6:mTurquoise2-NLS | AT1G66470 (*RHD6*) | *Arabidopsis thaliana* | pGGZ003 | This study |
| pGG-F-gATG5-G | AT5G17290 (*ATG5*) | *Escherichia coli* | pGGF000 | Adapted from<br>Decaestecker et al.<br>(2019) The Plant Cell. |
| ProEXP7:Cas9-GFP_gATG5 | AT1G12560 (*EXP7*);<br>AT5G17290 (*ATG5*) | *Arabidopsis thaliana* | pFASTR-AG | Adapted from<br>Decaestecker et al.<br>(2019) The Plant Cell. |
| ProRHD6:Cas9-GFP_gATG5 | AT1G66470 (*RHD6*);<br>AT5G17290 (*ATG5*) | *Arabidopsis thaliana* | pFASTR-AG | Adapted from<br>Decaestecker et al.<br>(2019) The Plant Cell. |
| pTwist-C-ATG5-D | AT5G17290 (*ATG5*) | *Escherichia coli* | pTwist Amp<br>High Copy | Twist Bioscience |
| ProEXP7:ATG5 | AT1G12560 (*EXP7*);<br>AT5G17290 (*ATG5*) | *Arabidopsis thaliana* | pGGZ003 | This study |
| ProRHD6:ATG5 | AT1G66470 (*RHD6*);<br>AT5G17290 (*ATG5*) | *Arabidopsis thaliana* | pGGZ003 | This study |

**Table 3.**
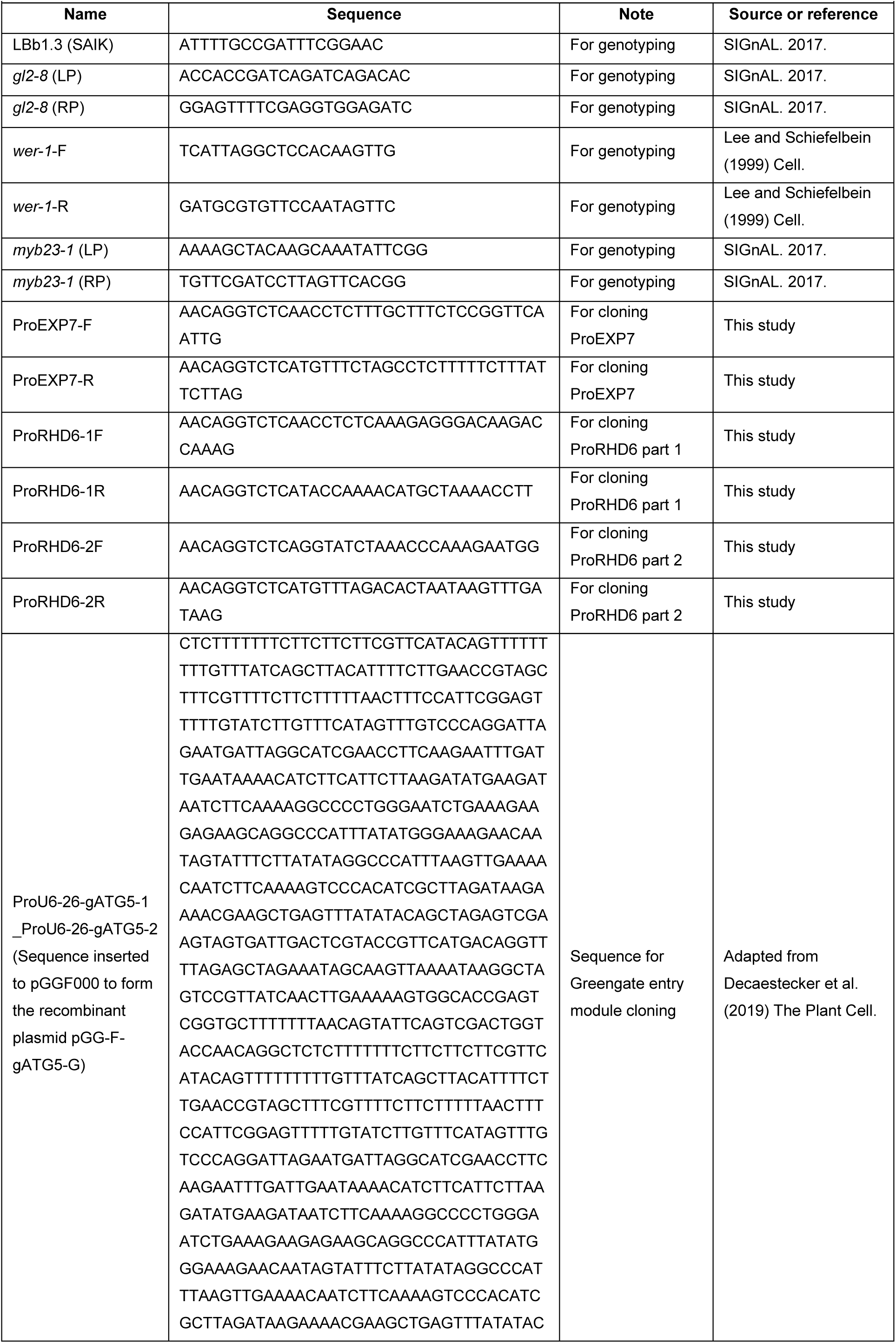
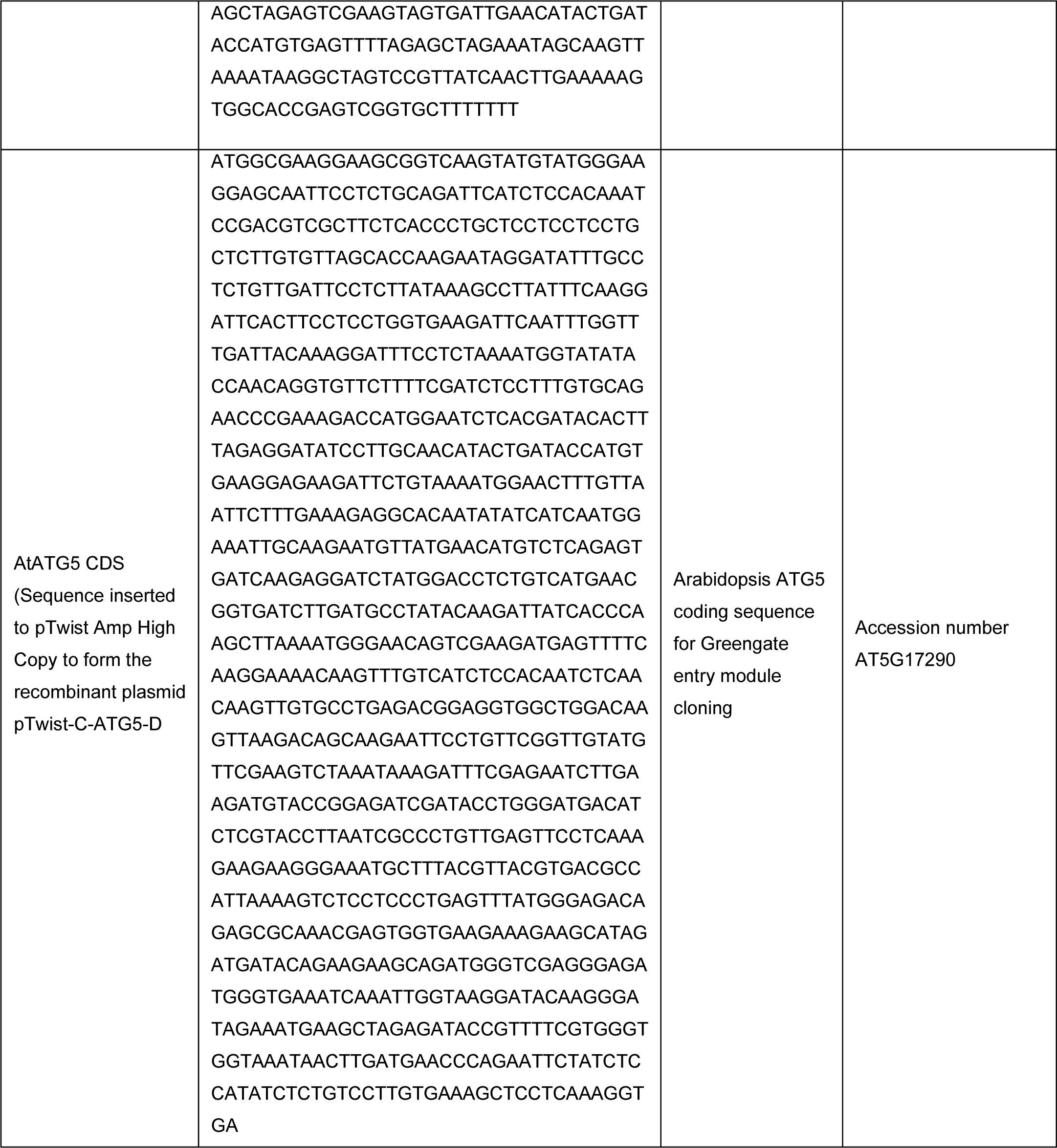
Primers and synthetic sequences used in this study.

All plasmids were assembled via the GreenGate cloning method (Lampropoulos et al., 2013). DNA sequences of *ProEXP7* and *ProRHD6* were cloned from Arbidopsis genome DNA using the primers listed in Table 3 and were subsequently ligated to pGGA000 to form the promoter entry modules pGG-A-ProEXP7-B and pGG-A-ProRHD6-B, respectively. These two modules and the promoter entry module pGG-A-ProUBQ10+Ω-B (Zhao et al., 2022) were further assembled with other GreenGate modules pGGB003 (Lampropoulos et al., 2013), pSW596-mTurquoise2 (Addgene plasmid # 115985; Schürholz et al., 2018), pGGD007 (Lampropoulos et al., 2013), pGGE009 (Lampropoulos et al., 2013), pGG-F-Alired-G (Zhao et al., 2022) and pGGZ003 (Lampropoulos et al., 2013) to form the plasmids ProUBQ10:mTurquoise2-NLS, ProRHD6:mTurquoise2-NLS and ProEXP7:mTurquoise2-NLS, respectively.

For constructing plasmids for CRISPR-TSKO, we used an adapted method from Decaestecker et al., 2019. The DNA fragments of two ProU6-26-driven guide RNAs for knocking out ATG5 (ProU6-26-gATG5-1_ProU6-26-gATG5-2; Table 3) were cloned and inserted into pGGF000 to form the entry module pGG-F-gATG5-G. This entry module, together with other GreenGate modules pGG-B-Linker-C, pGG-C-Cas9PTA*-D, pGG-D-P2A-GFP-NLS-E, pGG-E-G7T-F, pFASTR-A-G, was assmebled with either pGG-A-ProEXP7-B or pGG-A-ProRHD6-B to form the recombinant plasmids ProEXP7:Cas9-GFP_gATG5 and ProRHD6:Cas9-GFP_gATG5, respectively.

The coding sequence of Arabidopsis *ATG5* (Table 3) were cloned into pTwist Amp High Copy to form the entry module pTwist-C-ATG5-D (Twist Bioscience). The GreenGate modules pGGB003 (Lampropoulos et al., 2013), pTwist-C-ATG5-D (Twist Bioscience), pGGD002 (Lampropoulos et al., 2013), pGGE009 (Lampropoulos et al., 2013), pGG-F-Alired-G (Zhao et al., 2022) and pGGZ003 (Lampropoulos et al., 2013) were assembled with either pGG-A-ProEXP7-B or pGG-A-ProRHD6-B to form the recombinant plasmids ProEXP7:ATG5 and ProRHD6:ATG5, respectively.

All transgenic Arabidopsis lines were generated through the Agrobacterium-mediated floral-dip method (Clough and Bent, 1998). ProEXP7:Cas9-GFP_gATG5 and ProRHD6:Cas9-GFP_gATG5 were transformed into Arabidopsis wildtype Col-0 plants and the positive transformants were labeled as the line TSKO-E and TSKO-R, respectively. ProEXP7:ATG5 and ProRHD6:ATG5 were transformed into Arabidopsis *atg5-1* mutant plants and the positive transformants were labeled as the line E-ATG5 and R-ATG5, respectively.

### Plant phenotypic assays

Arabidopsis seeds were sterilized with 70% ethanol + 0.5‰ Tween 20 for 15 min and then with 100% ethanol for 15 min. Sterilized seeds were subsequently stored at 4°C for 2 d for vernalization. Vernalized seeds were sown 1/2 MS media (Murashige and Skoog salt + Gamborg B5 vitamin mixture [Duchefa] supplemented with 0.5 g/liter MES and 1% sucrose, pH 5.7) plates (+ 1% plant agar [Duchefa]) and vertically grown at 21°C at 60% humidity under LEDs with 70 mM/m^2^s a and a 16 h light/8 h dark photoperiod for 6 d. 6-d-old seedlings were subsequently transferred to 1/2 MS media plates (+1% plant agar [Duchefa]) containing 150 mM NaCl and vertically grown for 4 d. The seedlings were imaged at the day of transfer (d0) and at the fourth day after transfer (d4). The percentage of non-etiolated leaves to total leaves (including the cotyledons) of Arabidopsis seedlings at d4 were calculated for statistical analysis.

### Preparation of *Arabidopsis thaliana* samples for confocal microscopy

Arabidopsis seeds were were sterilized with 70% ethanol + 0.5‰ Tween 20 for 15 min and then with 100% ethanol for 15 min. Sterilized seeds were subsequently stored at 4°C for 2 d for vernalization. Vernalized seeds were spread on 1/2 MS media plates (+ 1% plant agar [Duchefa]) and vertically grown at 21°C at 60% humidity under LEDs with 50 mM/m^2^s a and a 16 h light/8 h dark photoperiod for 5 d. 5-day-old seedlings were placed in 1/2 MS media and treated with salt or chemicals as indicated in each experiment before confocal imaging. For nitrogen starvation, 1/2 MS media was replaced by nitrogen-deficient 1/2 MS media (Murashige and Skoog salt without nitrogen [CaissonLabs] + Gamborg B5 vitamin mixture [Duchefa] supplemented with 0.5 g/liter MES and 1% sucrose, pH 5.7).

For confocal microscopy, 5-d-old Arabidopsis seedlings were placed on a microscope slide with either water or water with 0.002 mg/ml propidium iodide and covered with a coverslip.

### Confocal microscopy

All images except the cotyledon epidermis imaging (Figure S7) were acquired by an upright point laser scanning confocal microscope ZEISS LSM800 Axio Imager.Z2 (Carl Zeiss) equipped with high-sensitive GaAsP detectors (Gallium Arsenide), a Plan-Apochromat 20X objective lens (numerical aperture 0.8, dry) and ZEN software (blue edition, Carl Zeiss). For cotyledon epidermis imaging (Figure S7), a LD C-Apochromat 40X objective lens (numerical aperture 1.1, water) was used instead of the 20X one. The GFP, mTurquoise2 and CoroNa Green AM fluorescence were excited at 488 nm and detected between 488 nm and 545 nm. The propidium iodide and mCherry fluorescence was excited at 561 nm and detected between 570 nm and 617 nm. For Z-stack imaging, interval between layers was set as 1 μm. Confocal images were processed with Fiji (version 1.54f, Fiji).

### Statistical analyses

All quantification analyses and statistical tests were performed with GraphPad Prism 8 software (version 8.1.1). For comparing the significance of differences between two experimental groups, paired and two-tailed student’s t-tests were performed as indicated in each experiment. For comparing the significance of differences between multiple experimental groups, Paired repeated measures one-way ANOVA and Fisher’s LSD tests were performed as indicated in each experiment.

## Acknowledgments

We thank George Haughn, Moritz Nowack, Ni Zhan and Qiangnan Feng for sharing experimental materials. We acknowledge funding from Austrian Academy of Sciences, Austrian Science Fund (FWF, P32355, P34944, I 6760, SFB F79, DOC 111), Vienna Science and Technology Fund (WWTF, LS17-047, LS21-009), European Research Council Grant (Project number: 101043370), and DFG-Heisenberg Professorship to Y.D. lab. We thank Vienna Biocenter Core Facilities (VBCF) Biooptics, Plant Sciences and Molecular Biology Service.

The authors declare no competing financial interests.

## Author contributions

J. Zhao designed the experiments, constructed plasmids, generated Arabidopsis lines, performed microscopy experiments and phenotypic assays, prepared figures, and wrote the draft. C. Löfke constructed plasmids, generated Arabidopsis lines and performed preliminary microscopy experiments. K. C. Yueng constructed plasmids and generated Arabidopsis lines. Y. Chen performed microscopy experiments and generated Arabidopsis lines. Y. Dagdas designed the experiments, supervised students and wrote the draft.

## Data availability

All the source data associated with the data presented in this manuscript is available at Zenodo. DOI: 10.5281/zenodo.15034967 and DOI: 10.5281/zenodo.15034948

---

**Figure S1.**
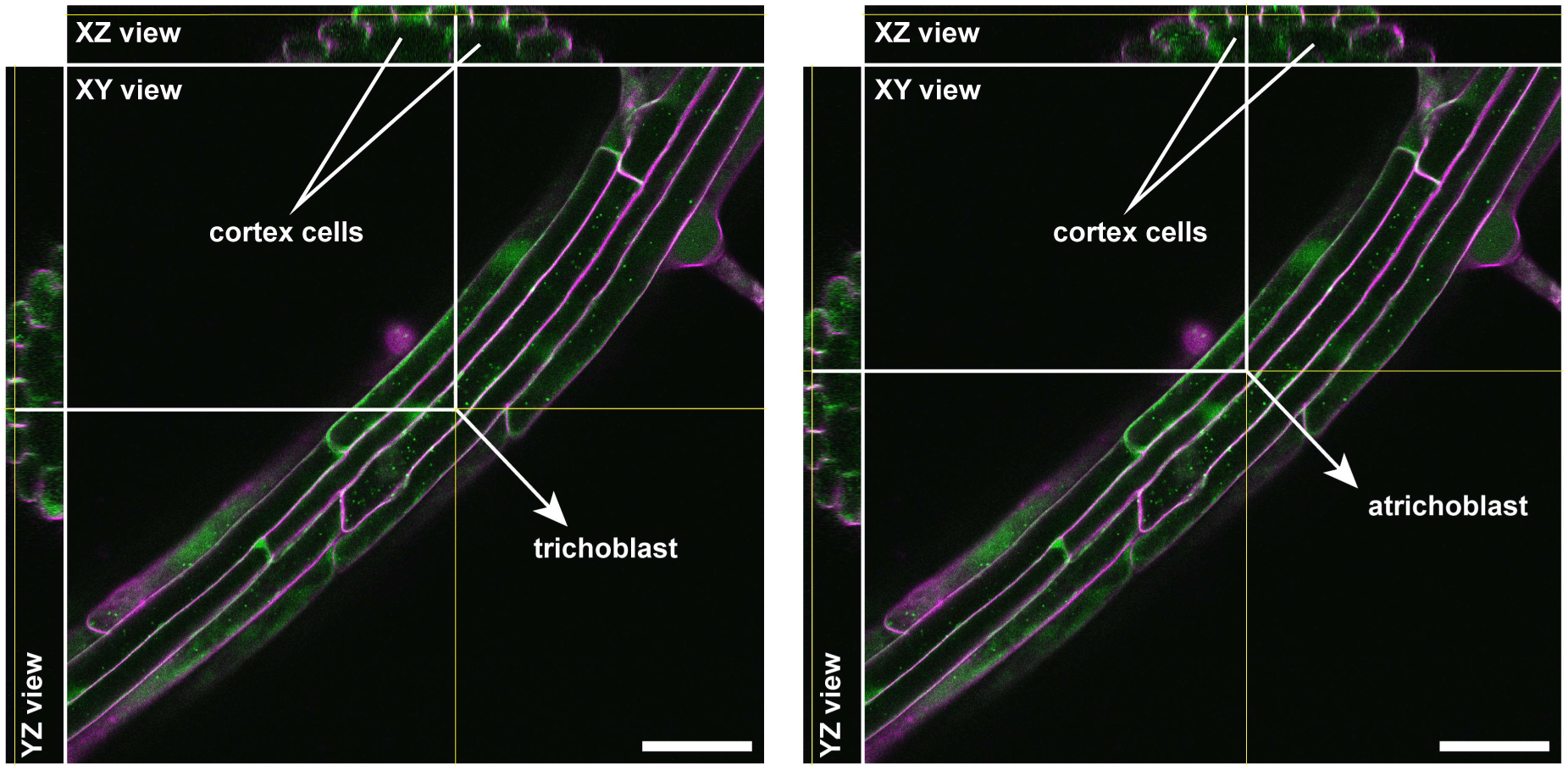
Schematic confocal images illustrating the approach used to distinguish trichoblasts from atrichoblasts through confocal microscopy. To identify trichoblast and atrichoblast cells, we analyzed XZ and YZ views of Z-stack images. Trichoblast (root hair forming) cells are identified as cells that are adjacent to two cortex cells and atrichoblast (non-root hair forming) cells as cells that are adjacent to one cortex cell. Green color, GFP-ATG8A. Magenta color, propidium iodide dye (for cell wall staining). Scale bars, 50 μm. Yellow lines show the alignment of XY, XZ and YZ views of the target cell.

**Figure S2.**
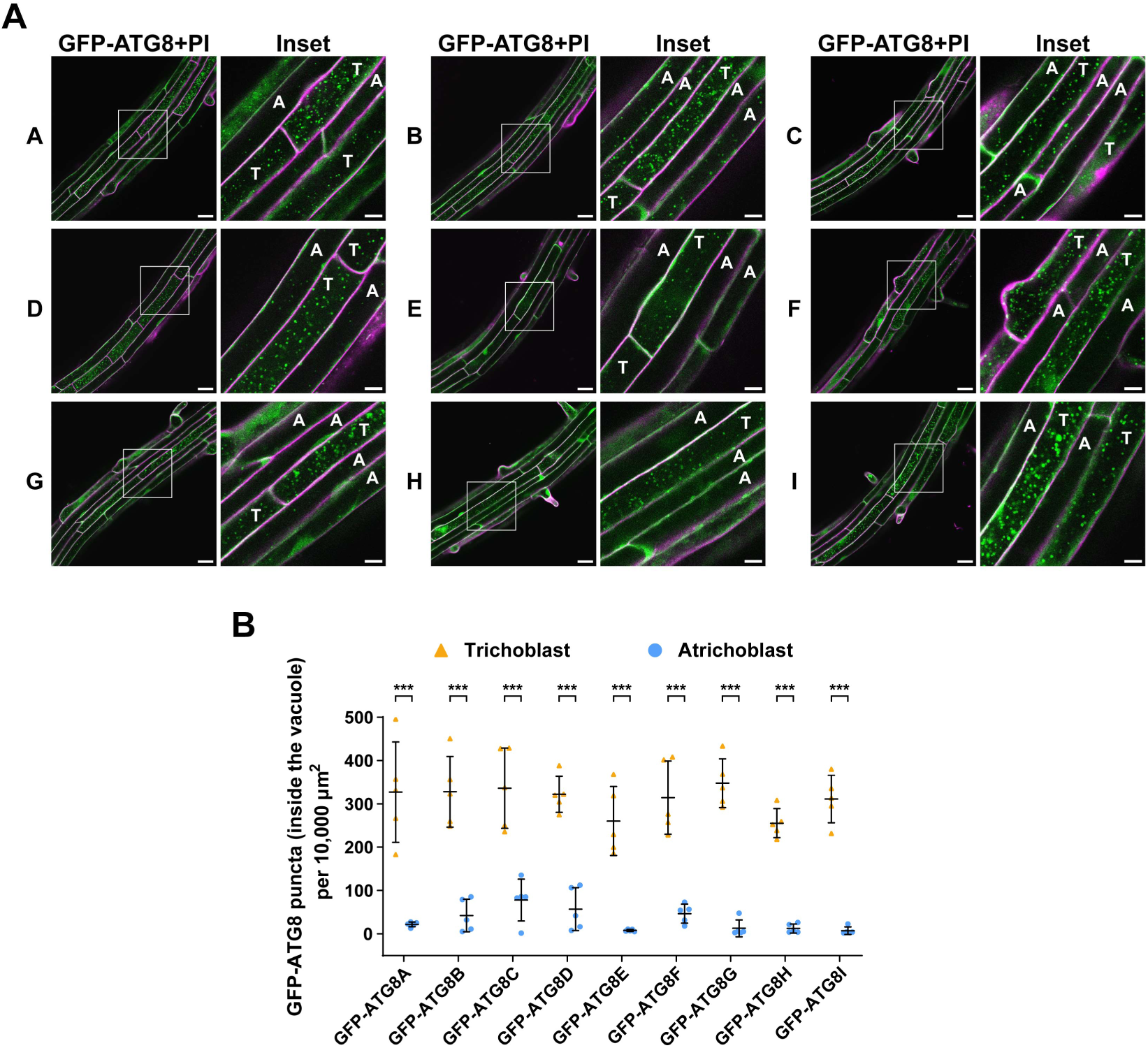
Trichoblasts exhibit higher autophagic flux than atrichoblasts under nitrogen-starvation conditions. **(A)** Confocal microscopy images of the trichoblasts and atrichoblasts at the root maturation zone of Arabidopsis wildtype Col-0 expressing ProUBQ10:GFP-ATG8A–I isoforms under nitrogen-starvation conditions. 5-d-old Arabidopsis seedlings were incubated in nitrogen-deficient 1/2 MS media containing 2 μm concanamycin A for 2 h before imaging. Representative images of 5 replicates are shown. Area highlighted in the white-boxed region in the GFP-ATG8+PI panel was further enlarged and presented in the inset panel. Scale bars, 30 μm. Inset scale bars, 10 μm. Green color, GFP-ATG8A–I isoforms. Magenta color, propidium iodide dye. T, trichoblasts. A, atrichoblasts. **(B)** Quantification of the GFP-ATG8 puncta inside the vacuole per normalized area (10,000 μm^2^) of the trichoblasts and atrichoblasts imaged in A. Bars indicate the mean ± SD of 5 replicates. Two-tailed and paired student t tests were performed to analyze the significance of GFP-ATG8 puncta density differences between the trichoblasts and the atrichoblasts. ***, P < 0.001

**Figure S3.**
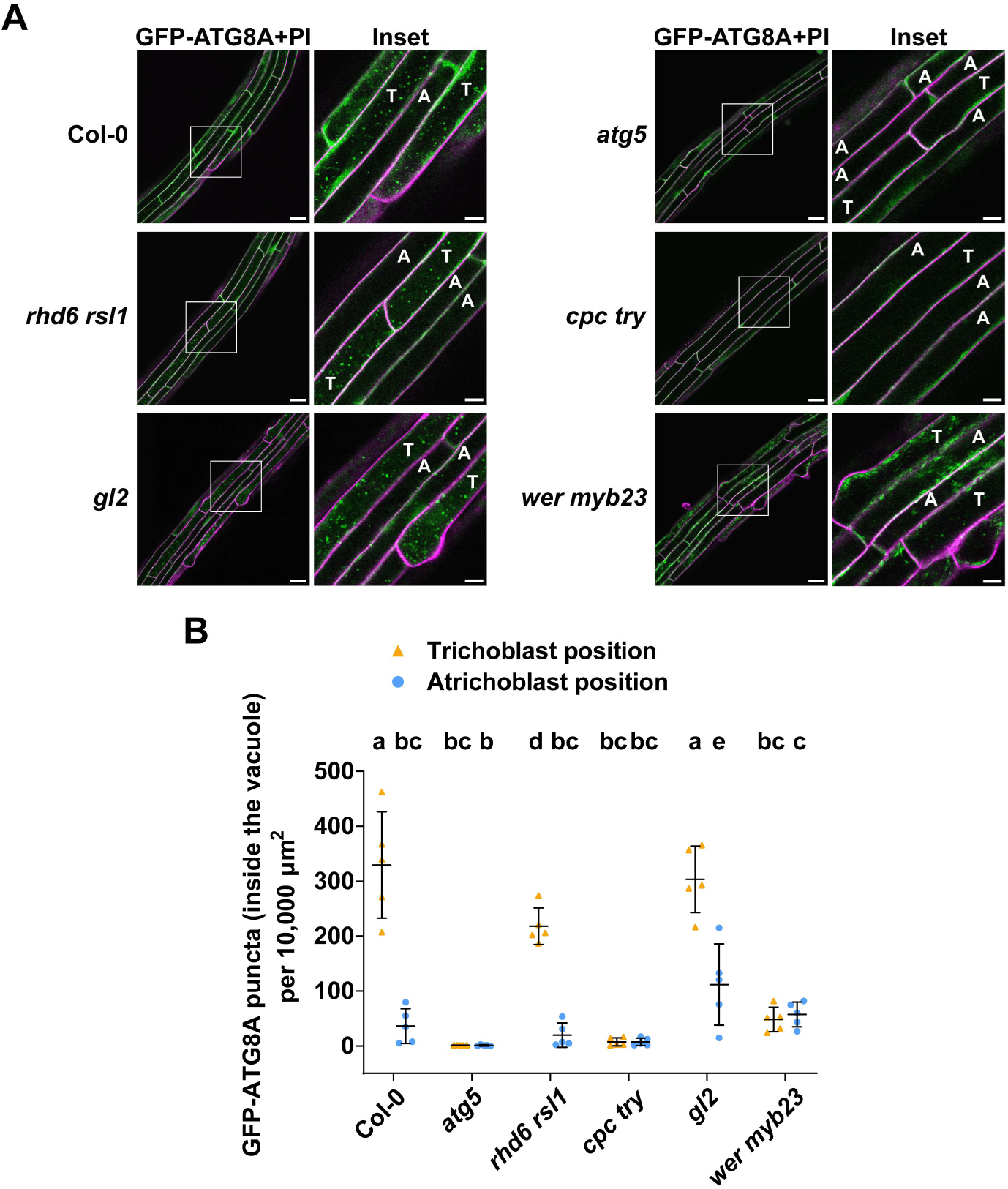
Genetic basis of the autophagic flux difference between trichoblasts and atrichoblasts. **(A)** Confocal microscopy images of epidermal cells at the root maturation zone of Col-0, *atg5*, *rhd6 rsl1*, *cpc try*, *gl2* and *wer myb23* expressing ProUBQ10:GFP-ATG8A under nitrogen-starvation conditions. 5-d-old Arabidopsis seedlings were incubated in nitrogen-deficient 1/2 MS media containing 2 μm concanamycin A for 2 h before imaging. Representative images of 5 replicates are shown. Area highlighted in the white-boxed region in the GFP-ATG8A+PI panel was further enlarged and presented in the inset panel. Scale bars, 30 μm. Inset scale bars, 10 μm. Green color, GFP-ATG8A. Magenta color, propidium iodide dye. Note, T indicates the trichoblast positions (adjacent to two cortex cells) and the A indicates the atrichoblast positions (adjacent to only one cortex cell), rather than the cell identities as mutants are affected in cell identity development. **(B)** Quantification of the GFP-ATG8A puncta inside the vacuole per normalized area (10,000 μm^2^) of the cells at the trichoblast positions and the atrichoblast positions imaged in A. Bars indicate the mean ± SD of 5 replicates. Paired repeated measures one-way ANOVA and Fisher’s LSD tests were used to analyze the differences of the number of the GFP-ATG8A puncta between each group. Family-wise significance and confidence level, 0.05 (95% confidence interval).

**Figure S4.**
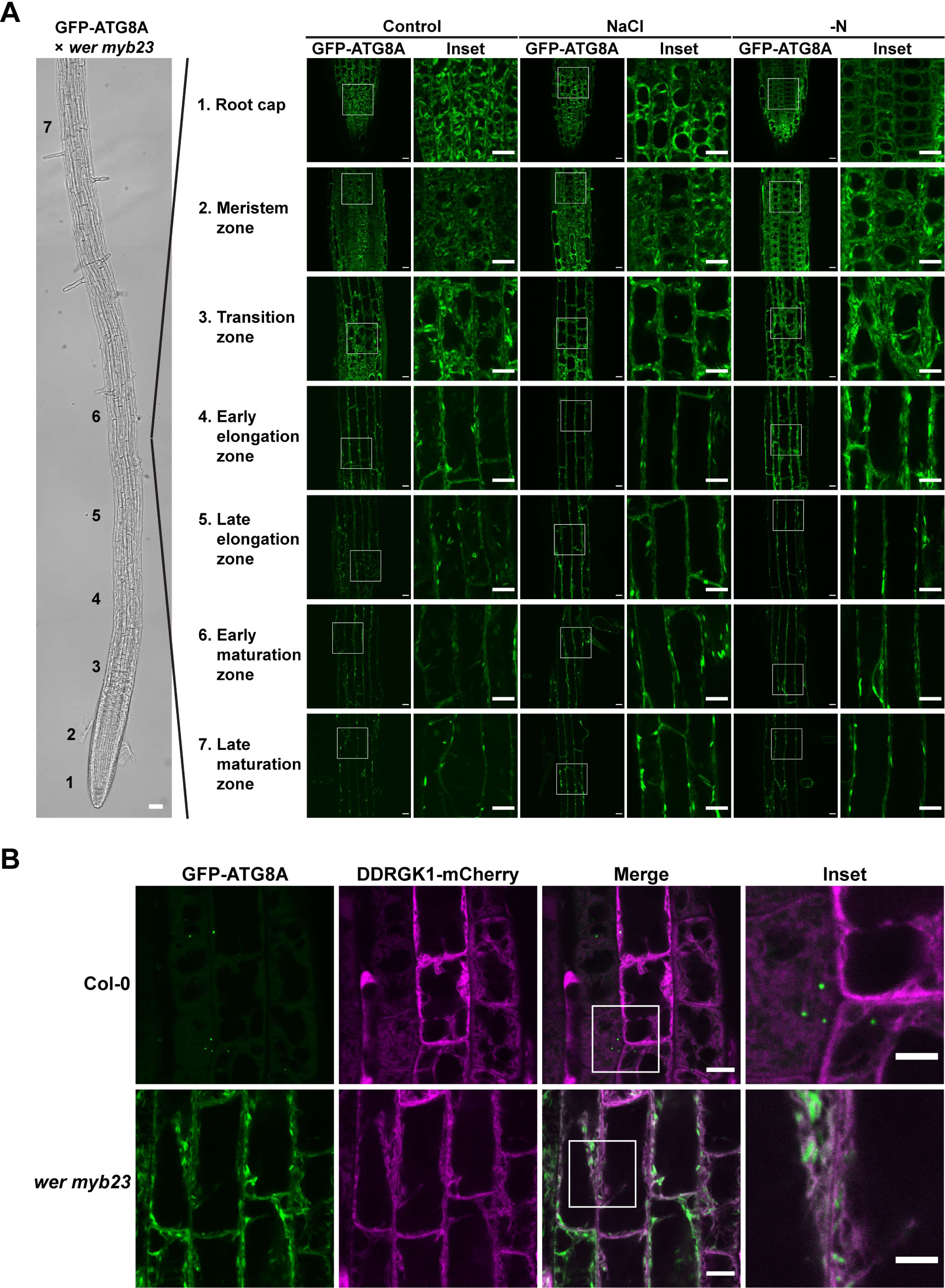
GFP-ATG8A is trapped at the endoplasmic reticulum in *wer myb23* mutant. **(A)** Confocal microscopy images of cells from different root regions of *wer myb23* expressing ProUBQ10:GFP-ATG8A. 5-d-old Arabidopsis seedlings were incubated in either control 1/2 MS media (Control) for 3 h, 100 mM NaCl-containing 1/2 MS media (NaCl) for 45 min, or nitrogen-deficient 1/2 MS media (−N) for 3 h before imaging. Representative images of 3 replicates are shown. Area highlighted in the white-boxed region in the GFP-ATG8A panel was further enlarged and presented in the inset panel. Bright field scale bars, 50 μm. GFP-ATG8A panel scale bars, 10 μm. Inset scale bars, 10 μm. **(B)** Confocal microscopy images of epidermal cells of the root transition zone of Arabidopsis wildtype Col-0 and *wer myb23* mutant lines co-expressing ProUBQ10:GFP-ATG8A and the endoplasmic reticulum marker ProUBQ10:DDRGK1-mCherry. 5-d-old Arabidopsis seedlings were incubated in 1/2 MS media for 3 h before imaging. Representative images of 2 replicates are shown. Area highlighted in the white-boxed region in the merge panel was further enlarged and presented in the inset panel. Scale bars, 10 μm. Inset scale bars, 5 μm.

**Figure S5.**
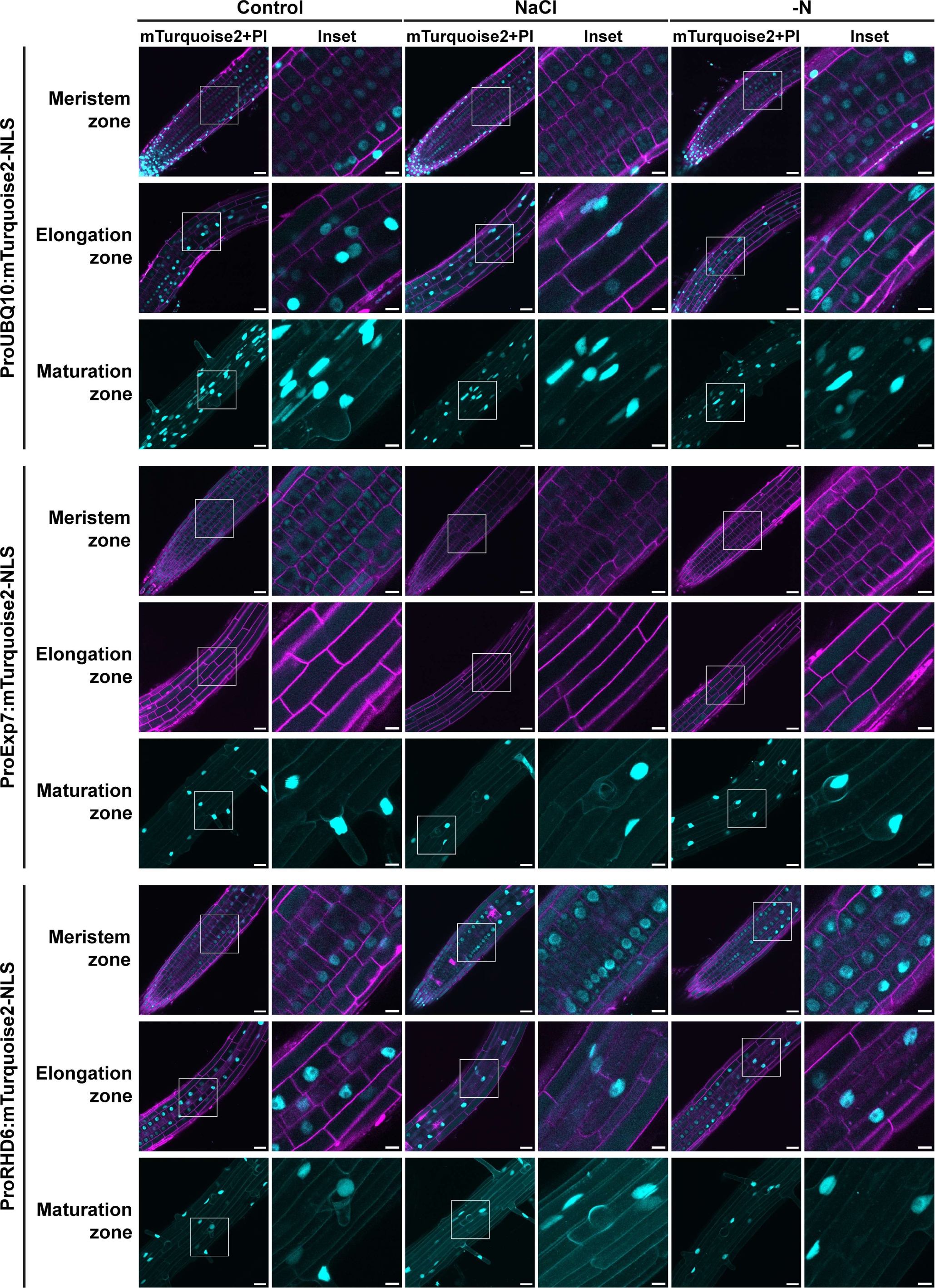
Expression patterns of *UBQ10*, *EXP7* and *RHD6* promoters in *Arabidopsis thaliana* roots. Confocal microscopy images of root epidermal cells of different root regions of Arabidopsis wildtype Col-0 expressing either ProUBQ10:mTurquoise2-NLS, ProEXP7:mTurquoise2-NLS or ProRHD6:mTurquoise2-NLS. 5-d-old Arabidopsis seedlings were incubated in either control 1/2 MS media (Control) for 3 h, 100 mM NaCl-containing 1/2 MS media (NaCl) for 45 min, or nitrogen-deficient 1/2 MS media (−N) for 3 h before imaging. For images showing the maturation zone, an mTurquoise2-single channel Z-stack image was shown. Representative images of 3 replicates are shown. Area highlighted in the white-boxed region in the mTurquoise2+PI panel was further enlarged and presented in the inset panel. Scale bars, 30 μm. Inset scale bars, 10 μm. Cyan color, mTurquoise2-NLS. Magenta color, propidium iodide dye. NLS, nuclear location signal peptide.

**Figure S6.**
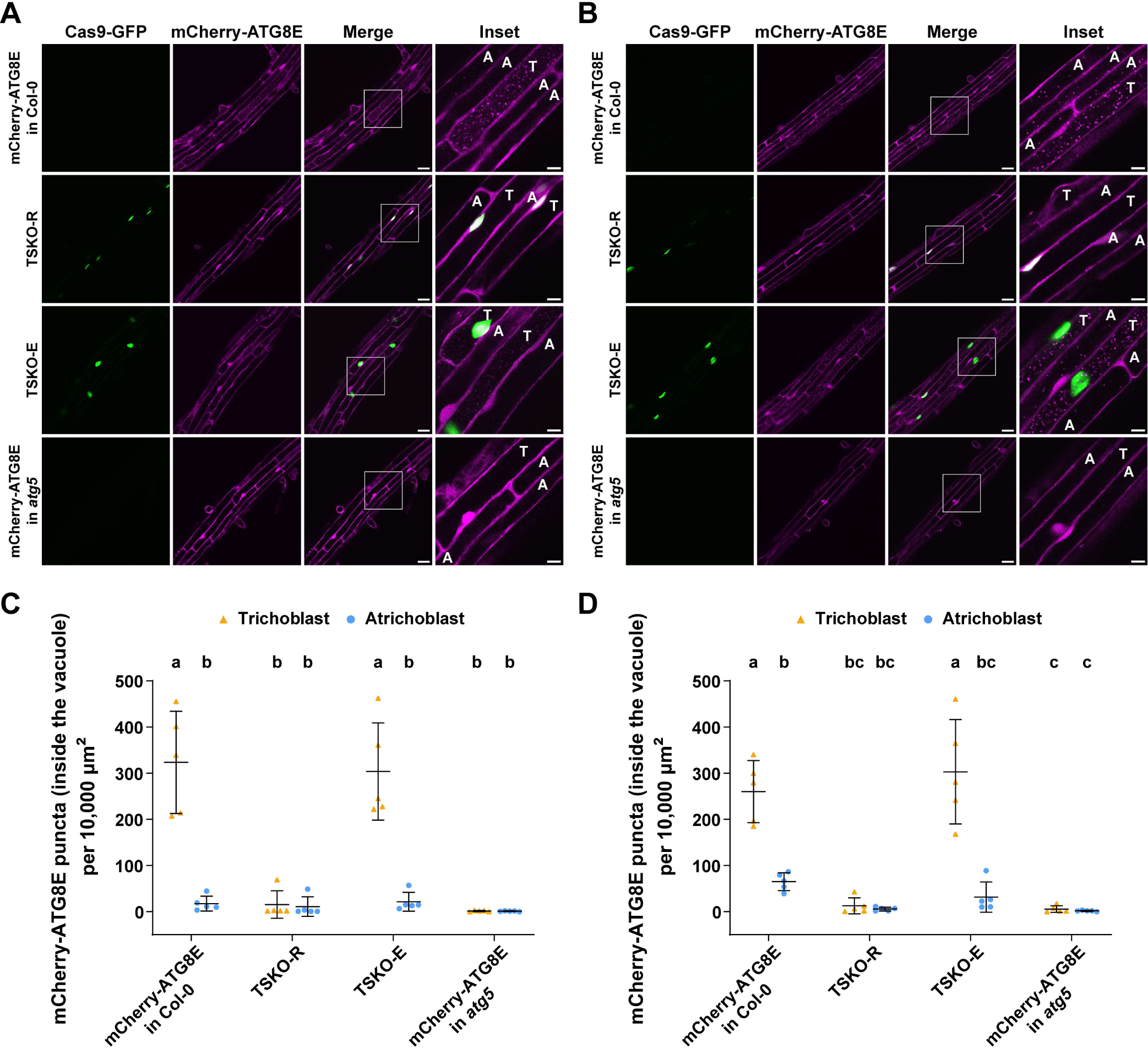
*RHD6* promoter was more efficient in disrupting autophagic flux in trichoblast cells compared to *EXP7* promoter in tissue specific CRISPR mutagenesis experiments. **(A)** Confocal microscopy images of trichoblasts and atrichoblasts at the root maturation zone of Arabidopsis lines mCherry-ATG8E in Col-0, TSKO-R (*ATG5* mutagenized using *ProRHD6*-driven Cas9), TSKO-E (*ATG5* mutagenized using *ProEXP7*-driven Cas9) and mCherry-ATG8E in atg5 under control treatment. 5-d-old Arabidopsis seedlings were incubated in 1/2 MS media containing 2 μm concanamycin A for 2 h before imaging. Representative images of 5 replicates are shown. Area highlighted in the white-boxed region in the merge panel was further enlarged and presented in the inset panel. Scale bars, 30 μm. Inset scale bars, 10 μm. T, trichoblasts. A, atrichoblasts. **(B)** Confocal microscopy images of trichoblasts and atrichoblasts at the root maturation zone of Arabidopsis lines mCherry-ATG8E in Col-0, TSKO-R (*ATG5* mutagenized using *ProRHD6*-driven Cas9), TSKO-E (*ATG5* mutagenized using *ProEXP7*-driven Cas9) and mCherry-ATG8E in atg5 under NaCl stress treatment. 5-d-old Arabidopsis seedlings were incubated in 1/2 MS media containing 50 mM NaCl + 1 μm concanamycin A for 1 h before imaging. Representative images of 5 replicates are shown. Area highlighted in the white-boxed region in the merge panel was further enlarged and presented in the inset panel. Scale bars, 30 μm. Inset scale bars, 10 μm. T, trichoblasts. A, atrichoblasts. **(C)** Quantification of the mCherry-ATG8E puncta inside the vacuole per normalized area (10,000 μm^2^) of the cells of the trichoblasts and atrichoblasts imaged in A. Bars indicate the mean ± SD of 5 replicates. Paired repeated measures one-way ANOVA and Fisher’s LSD tests were used to analyze the differences of the number of the mCherry-ATG8E puncta between each group. Family-wise significance and confidence level, 0.05 (95% confidence interval). **(D)** Quantification of the mCherry-ATG8E puncta inside the vacuole per normalized area (10,000 μm^2^) of the cells of the trichoblasts and atrichoblasts imaged in B. Bars indicate the mean ± SD of 5 replicates. Paired repeated measures one-way ANOVA and Fisher’s LSD tests were used to analyze the differences of the number of the mCherry-ATG8E puncta between each group. Family-wise significance and confidence level, 0.05 (95% confidence interval).

**Figure S7.**
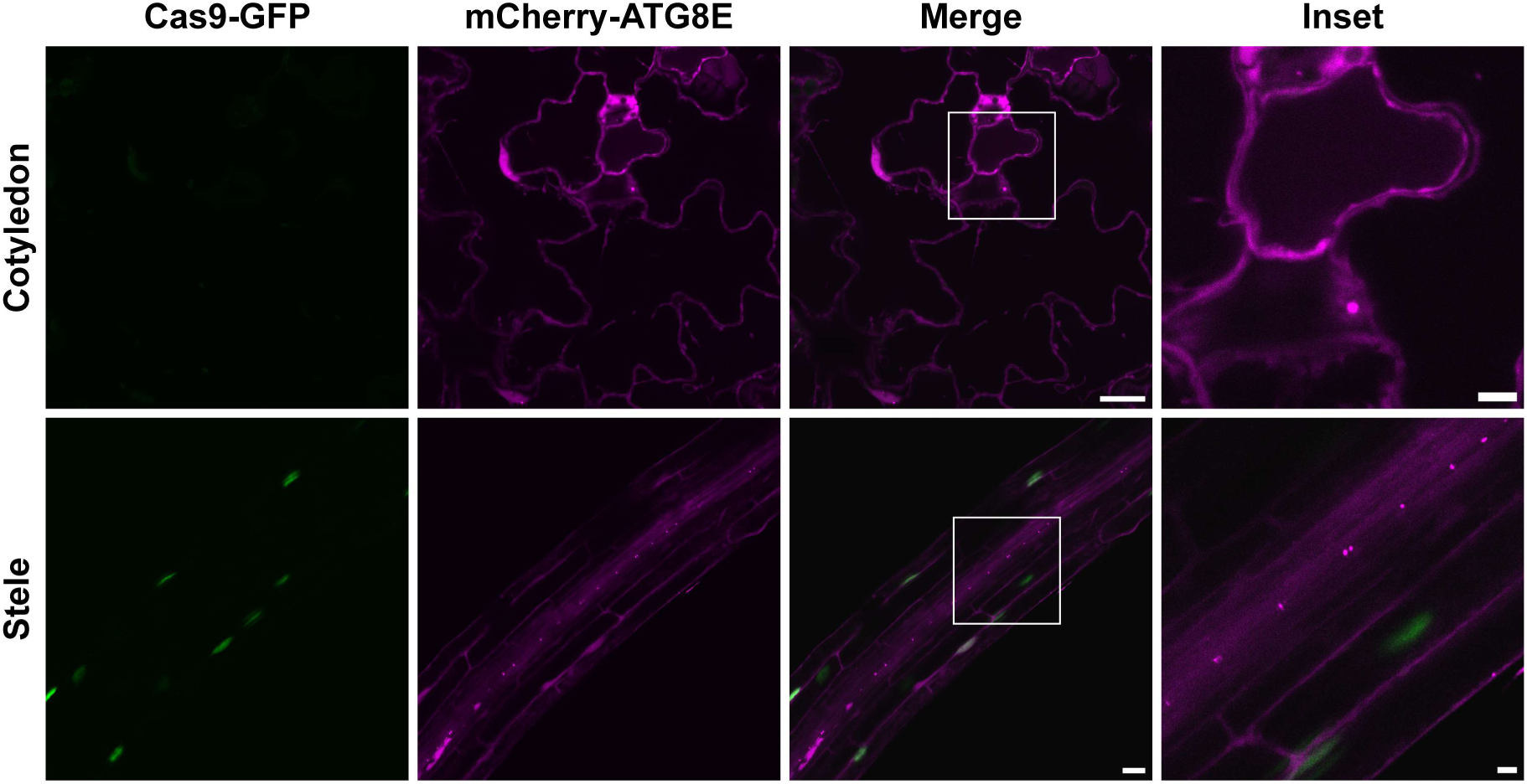
Normal autophagosome structures can be observed in cells that donot express Cas9 in TSKO-R lines. Confocal microscopy images of cells of cotyledon epidermis and stele regions of TSKO-R (*ATG5* mutagenized using *ProRHD6*-driven Cas9). 5-d-old Arabidopsis seedlings were incubated in control 1/2 MS media for 30 min before imaging. Representative images of 3 replicates are shown. Area highlighted in the white-boxed region in the merge panel was further enlarged and presented in the inset panel. Scale bars, 20 μm. Inset scale bars, 5 μm.

